# Domain-adaptive matching bridges synthetic and *in vivo* neural dynamics for neural circuit connectivity inference

**DOI:** 10.1101/2022.10.03.510694

**Authors:** Kaiwen Sheng, Shanghang Zhang, Shenjian Zhang, Yutao He, Maxime Beau, Peng Qu, Xiaofei Liu, Youhui Zhang, Lei Ma, Kai Du

**Affiliations:** Institute for Artificial Intelligence, Peking University, Beijing, China; School of Computer Science, Peking University, Beijing, China; Beijing Academy of Artificial Intelligence, Beijing, China; Department of Computer Science and Technology, Tsinghua University, Beijing, China; Department of Psychological and Cognitive Sciences, Tsinghua University, Beijing, China; Wolfson Institute for Biomedical Research, University College London, London, United Kingdom

**Keywords:** *in-vivo* recording, extracellular electrophysiology, neural circuit, synaptic connectivity, deep learning, model mismatch, out-of-distribution (OOD)

## Abstract

Accurately inferring neural circuit connectivity from *in vivo* recordings is essential for understanding the computations that support behavior and cognition. However, current deep learning approaches are limited by incomplete observability and the lack of ground-truth labels in real experiments. Consequently, models are often trained on synthetic data, which leads to the well-known “model mismatch” problem when simulated dynamics diverge from true neural activity. To overcome these challenges, we present Deep Domain-Adaptive Matching (DeepDAM), a training framework that adaptively *matches* synthetic and *in vivo* data domains for neural connectivity inference. Specifically, DeepDAM fine-tunes deep neural networks on a combined dataset of synthetic simulations and unlabeled *in vivo* recordings, aligning the model’s feature representations with real neural dynamics to mitigate model mismatch. We demonstrate this approach in rodent hippocampal CA1 circuits as a proof-of-concept, achieving near-perfect connectivity inference performance (Matthews correlation coefficient ∼0.97–1.0) and substantially surpassing classical methods (∼0.6–0.7). We further demonstrate robustness across multiple recording conditions within this hippocampal dataset. Additionally, to illustrate its broader applicability, we extend the framework to two distinct systems without altering the core methodology: a stomatogastric microcircuit in *Cancer borealis* (*ex vivo*) and single-neuron intracellular recordings in mouse, where DeepDAM significantly improves efficiency and accuracy over standard approaches. By effectively leveraging synthetic data for *in vivo* and *ex vivo* analysis, DeepDAM offers a generalizable strategy for overcoming model mismatch and represents a critical step towards data-driven reconstruction of functional neural circuits.

## Introduction

Neural circuit connectivity underlies information processing and transmission within the brain, fundamentally shaping network dynamics, behavior, and cognition (1–4). Therefore, accurately reconstructing this connectivity from *in vivo* recordings is essential for deciphering the computational mechanisms that enable brain function. However, the intrinsic complexity and heterogeneity of neural networks—characterized by diverse neuron types and intricate synaptic interactions—pose serious challenges for connectivity inference. A primary issue is that even state-of-the-art *in vivo* recording techniques (5–7) capture only a limited subset of neurons within a vast network, leading to incomplete or ambiguous observations. Moreover, researchers often lack detailed prior knowledge of the specific circuitry under investigation, further complicating inference.

One strategy to address these incomplete observations is *model-based inference*, which relies on a hypothesized model to simulate (or synthesize) neural activity comparable to experimental data and then infers the connections that best explain the observations (8). Traditionally, unsupervised model-based methods such as the inverse Ising model (9) and Generalized Linear Models (GLMs) (10, 11) have been widely used for this purpose. These approaches can be effective in simpler neural circuits (for example, in early sensory areas); however, they struggle in brain regions with dense recurrent connectivity. In fact, Das and Fiete (8) showed using synthetic benchmarks that such methods face an inevitable limitation in complex networks with strong recurrent loops. This shortcoming—referred to as the *“model mismatch”* problem—arises when the simplistic or assumed dynamics of the computational model diverge from the true neural dynamics, causing inference errors.

In recent years, deep learning has emerged as a powerful alternative for neural connectivity inference, owing to its capacity to capture complex patterns in large datasets. Deep learning methods typically train a deep neural network (DNN) entirely on synthetic data generated from a chosen biophysical model (12), thereby sidestepping the need for exhaustive ground-truth labels (which are practically unattainable for *in vivo* recordings). Indeed, our preliminary investigations indicated that a DNN with a ResNet-LSTM architecture outperforms traditional model-based approaches on purely synthetic datasets (13). However, when we applied such a model—trained on synthetic data—to real *in vivo* recordings, its performance degraded markedly. This gap in performance prompted us to scrutinize the root cause: the disparity between synthetic training data and real neural data.

In this work, we identified a substantial difference between the feature distributions of synthetic spike data and those of real *in vivo* recordings, underscoring the severity of the model mismatch problem. This insight led us to develop Deep Domain-Adaptive Matching (DeepDAM) — a model-based deep learning framework tailored for neural connectivity inference that leverages both unlabeled *in vivo* spike trains and synthetic neural data. The key innovation of DeepDAM is to reframe the classical model mismatch issue as a data distribution mismatch, essentially treating it as an out-of-distribution (OOD) challenge. While domain adaptation for OOD data is well-established in machine learning (14, 15), to our knowledge it has not been applied in this particular context within computational neuroscience.

The DeepDAM framework operates in two stages: first by employing domain adaptation to align the DNN’s feature space with the dynamics of real neural data (14, 15), and then by using self-training to iteratively refine the network on unlabeled *in vivo* data (16–20). This two-phase strategy mitigates discrepancies between the synthetic and real data domains, thereby improving inference accuracy when the model is applied to real experiments. Since *in vivo* datasets with definitive connectivity labels are exceedingly scarce, validating our approach on a rigorously labeled *in vivo* dataset is a notable aspect of this study. To our best knowledge, we are the first to leverage this particular labeled dataset for model validation, and among the first to explicitly address model mismatch from an OOD perspective in neural circuit inference.

Using this unique dataset, we demonstrate that DeepDAM can accurately infer monosynaptic connections from multi-electrode extracellular recordings in the mouse hippocampal CA1 region. Notably, CA1 is a challenging testbed due to limited prior circuit knowledge and strongly recurrent connections that introduce additional confounds (8) (Fig. 1a). By coupling a ResNet-LSTM architecture with our domain-adaptive training strategy, the framework achieved near-perfect connectivity inference performance, with Matthews correlation coefficient (MCC) values around 0.97–1.0. This is a dramatic improvement over popular model-based inference methods (such as GLMs and other statistical models) (10, 12), which typically reach only ∼0.6–0.7 on the same task. Moreover, DeepDAM maintained high accuracy across multiple recording sessions and experimental conditions, each presenting varying levels of data ambiguity. This robustness under different noise levels and experimental perturbations highlights the framework’s reliability and adaptive capacity in realistic, less-controlled scenarios.

**Fig. 1.**
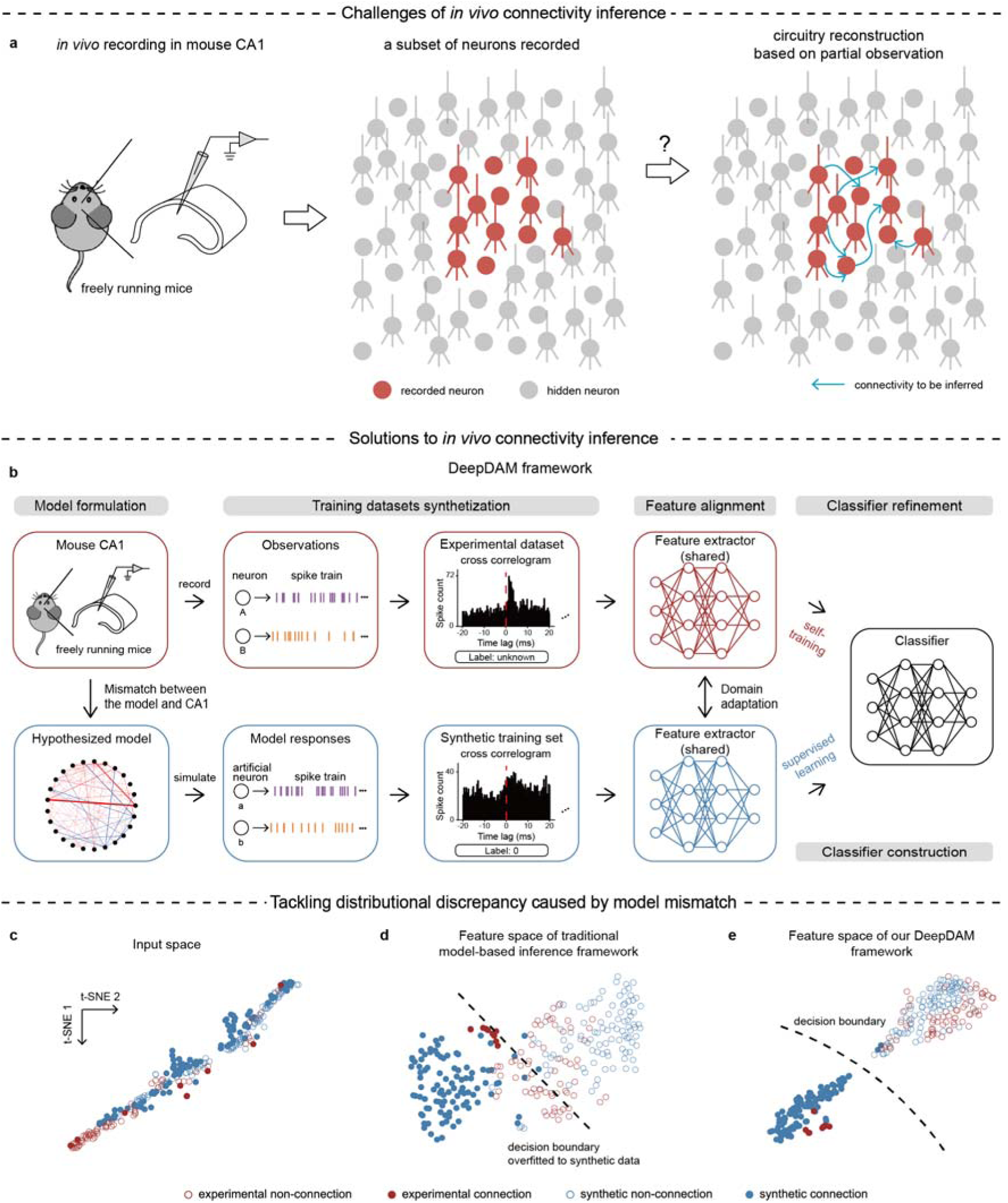
DeepDAM inference framework for monosynaptic connectivity inference in CA1 networks. (**a**) Challenges associated with inferring connectivity, including incomplete observations and strong recurrent connections. (**b**) DeepDAM framework. The gray boxes outlined the five major steps of the framework. Note that both the feature extractor and the classifier are shared between synthetic and experimental data. (**c**) Distributional discrepancy of experimental and synthetic data caused by model mismatch in the model construction step. (**d**) Synthetic and experimental feature distributions after training the traitional model-based inference framework. (**e**) Synthetic and experimental feature distributions after training our DeepDAM framework.

Because the hippocampal dataset is, to our knowledge, the only *in vivo* resource with both spike recordings and ground-truth connectivity, we also explored DeepDAM’s generalization to other neural inference problems across different species and scales. In these proof-of-concept explorations, we applied the framework to classic system identification tasks ranging from single neurons to small circuits. For example, DeepDAM successfully recovered biophysical parameters from large-scale intracellular recordings of individual mouse neurons (21, 22), and it identified synaptic connectivity in a well-characterized microcircuit (the stomatogastric ganglion of *Cancer borealis*). In each case, our approach outperformed standard model-fitting techniques (23) in both accuracy and efficiency, suggesting that the underlying domain-adaptive strategy has broad potential applicability beyond the initial hippocampal circuit demonstration.

In summary, DeepDAM offers a new paradigm for neural circuit connectivity inference by seamlessly integrating synthetic model data with real *in vivo* data through domain adaptation. Rather than a one-off use of domain adaptation or self-training, it provides a general framework that mitigates model mismatch and extends the power of computational models to more faithfully capture real neural dynamics. We emphasize that our study is presented as a proof-of-concept in rodent hippocampal circuits. Nonetheless, this demonstration illustrates the potential of DeepDAM to be extended to other brain regions and species in future work. By addressing the long-standing model mismatch problem, our approach paves the way for more accurate data-driven reconstructions of neural circuits. Ultimately, we envision that such domain-adaptive techniques will contribute to the ambitious goal of reconstructing mammalian brain connectivity from large-scale *in vivo* data.

## Results

### Addressing model mismatch in neural connectivity inference as an out-of-distribution problem

There are two primary challenges arising in the efforts of inferring monosynaptic connectivity from *in vivo* spike trains: (1) Incomplete observations, where the interaction between the recorded and unrecorded neurons can falsely enhance spike-timing correlations, leading to potential misinterpretations (**Fig. 1a**); and (2) Unknown network complexity, where computational models cannot fully cover the intricate nature of brain networks, such as cell-type-specific biophysics and recurrent connections. These challenges manifest themselves as significant ‘model mismatch’, i.e., discrepancies between the response of the computational model and the actual dynamics of the neural network (as shown in **Fig. 1c**). We further highlight this mismatch through t-distributed stochastic neighbor embedding (t-SNE) visualizations (24), revealing a clear misalignment between model-generated cross-correlograms (CCGs) from synthetic data and those directly obtained from the extracellular multi-electrode recordings in CA1 (25) when displayed in a two-dimensional space (**Fig. 1c**).

The basic idea of solving the model mismatch problem is to consider it as an OOD problem in machine learning, i.e., the performance is significantly degraded due to the mismatch in distribution between the model-generated dynamics and the actual neural dynamics. Domain adaptation techniques are commonly used to deal with the OOD problem by effectively bridging the gap between different data distributions (26). Based on recent advances in unsupervised domain adaptation techniques, instead of narrowing such gap in the original data space, we focus on the shared and informative features of synthetic and experimental data (i.e., feature space) (14, 27, 28). Accordingly, we developed a comprehensive, DNN-based framework for inferring monosynaptic connectivity from unlabeled *in vivo* extracellular recordings:

1. **Model formulation**: A multiple-timescale adaptive threshold (MAT) network (**Fig. 1b**), is formulated to mimic the dynamics of the underlying neural network. This MAT network is a recurrent neural network with excitatory and inhibitory synaptic weights that follow lognormal distributions (**Fig. S1b**) to generate firing rate distributions similar to hippocampal neural recordings (**Fig. S1a**) (12).
2. **Training dataset synthetization**: Exploiting the transparency of this computational model, a synthetic dataset is produced, providing ground-truth labels for cross-correlograms (CCGs) of neuron pairs (**Fig. 1b**). These CCGs, a series of time-lagged correlations between two spike trains, serve as crucial tools for understanding neural connectivity patterns (10, 12, 25, 29, 30).
3. **Feature alignment**: Features that are shared between the synthetic CCGs and experimental CCGs and informative of the underlying connectivity are extracted from DNN-based feature extractors using domain adaptation (**Fig. 1b**).
4. **Classifier construction**: The DNN’s classifier is first trained on labeled synthetic data via supervised learning. This step complements feature alignment by making sure that the features are informative about neural connectivity (**Fig. 1b**).
5. **Classifier refinement**: To address potential overfitting of the DNN’s classifier to synthetic data, we refine the classifier through self-training using unlabeled experimental data. This is achieved by assigning pseudo labels to the experimental data (**Fig. 1b**).

Our primary contribution to this field is the transformation of the conventional three-step model-based inference method (comprising steps 1, 2, and 4) into a five-step, DNN-based framework (‘DeepDAM’). Traditional inference methods, lacking feature alignment and classifier refinement steps, are prone to training biases, as both the feature extractor and classifier are trained exclusively on labeled synthetic data, ignoring the experimental dataset. This results in the classifier overfitting to synthetic features and skewing inferences when applied to experimental data (**Fig. 1d**, **Fig. S2a** & **b**). In contrast, our construction of DeepDAM, which includes Feature alignment and Classifier refinement, can effectively address these shortcomings, thereby minimizing the gap in feature space, preventing the classifiers from overfitting the synthetic data, and ensuring more accurate inferences on experimental data (**Fig. 1e** & **Fig. S2c**).

It is noteworthy that, unlike traditional model-fitting methods, which typically focus on adjusting model parameters to fit data, our approach emphasizes ‘model-generation’, where the MAT model is used as a generative tool to create synthetic training datasets without fine-tuning the MAT model parameters. This shifts from direct model optimization to ‘model-generation’ is crucial, as fine-tuning the biophysical network model is often impractical due to the complexities of the underlying biological mechanisms and the extensive number of model parameters (see Discussions).

In the following sections, we will first elaborate on key design elements of the DeepDAM framework’s implementation, highlighting its inherent flexibility for diverse applications. Next, we will further validate the performance and robustness of the DeepDAM in comparison to traditional inference methods, using real *in vivo* spike data for inferring monosynaptic connectivity. Finally, we will illustrate the versatility of DeepDAM framework by extending its application to traditional model-fitting tasks, such as inferring biophysical properties of large-scale single neurons from *in vitro* and a microcircuit from *ex vivo* data.

### Key implementations of DeepDAM

In our DeepDAM framework, we strategically incorporate and customize two core machine learning strategies—domain adaptation and self-training—to address the OOD challenges inherent in inferring monosynaptic connectivity from *in vivo* recordings. This fusion involves several critical designs, which we will elaborate on below.

The first key design element of DeepDAM is the formulation of composite loss functions for the DNN. In addition to the standard loss function for assessing regular connectivity accuracy, we introduce a composite loss function *L* comprising the domain adaptation loss (*L_DA_*) and the self-training loss (*L_ST_*) given by Equation 1. Here, *L_DA_* facilitates Feature Alignment, while *L_ST_* is crucial for Classifier Refinement.

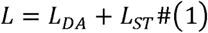

The domain adaptation loss in our DeepDAM framework is derived from unsupervised domain adaptation techniques (14, 27, 28) to reduce the distributional discrepancy between synthetic data (*x_syn_*) and unlabeled experimental data (*x_exp_*) in the feature space, as indicated in Equation 2 and 3:

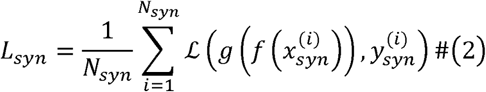

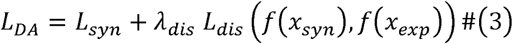

where 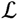 is the cross-entropy loss function 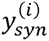 is the label for 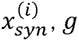 is the classifier and *λ_dis_* is the scaling factor of the discrepancy loss. Next, the synthetic and experimental features are categorized into clusters through the spherical K-means clustering algorithm, based on synthetic labels and pairwise feature similarity (Methods) (14). Our aim is to strategically minimize the distance *L_dis_* between features within the same cluster and maximize it between different clusters (**Fig. 2b**; Methods). At last, the feature extractor *f* and the classifier *g* are co-trained via the DNN to ensure high performance on labeled synthetic data. Overall, the domain adaptation loss (*L_DA_*) facilitates the transfer of this high performance from synthetic to experimental data.

**Fig. 2.**
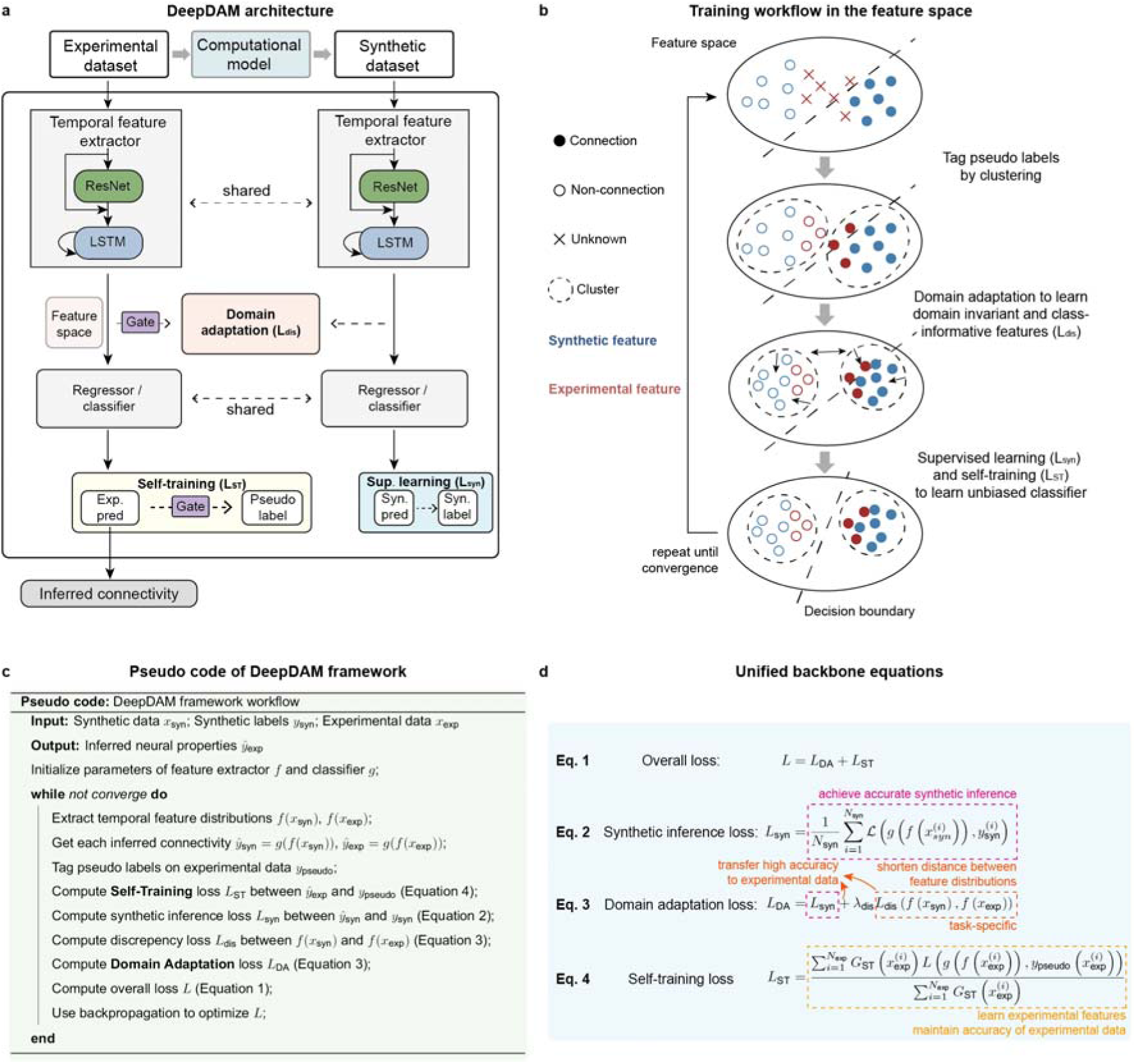
DeepDAM framework architecture. (**a**) Workflow of the DeepDAM framework. The temporal feature extractor and regressor/classifier are shared between synthetic data and experimental data. Firstly, temporal features of synthetic data and experimental data are extracted by the shared temporal feature extractor. The output space from the temporal feature extractor is the feature space. Secondly, the shared regressor or classifier processes the temporal features and outputs the inferred neural properties. Sup. learning stands for supervised learning. (**b**) The training workflow of our inference framework. Each dot or cross represented a sample feature. **(c)** Pseudo code. The equation IDs correspond to the IDs in the main text and **d**. **(d)** The function of each unified backbone equation.

However, even with domain adaptation, the classifier is not optimized on experimental features and the feature alignment process cannot perfectly align these two feature distributions, potentially making the classifier overfit on synthetic features. Therefore, we integrated self-training to explicitly refine the classifier to the experimental features (**Fig. 2b**) (16–19). Self-training applies “supervised learning” on unlabeled experimental data by tagging pseudo-labels to them, which are given by the synthetic data labels within the same cluster (**Fig. 2b** & **d**; Equation 4).

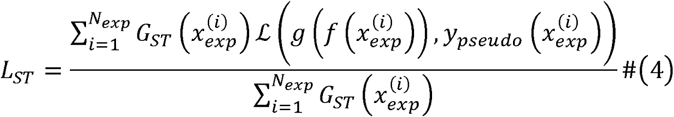

where 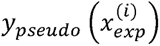 is the pseudo label of experimental datum 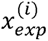 and *G* (·) is a binary function for gating 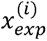 to the self-training process.

A second key design element in our DeepDAM framework is the introduction of a gating mechanism. At the beginning of the training process, the features extracted and the pseudo-labels assigned may be noisy and inaccurate. To address this, we introduce a gating mechanism that filters out experimental data if their distances to the designated cluster center exceed a predefined threshold. This filtration occurs during the domain adaptation and self-training phases of the current training iteration (as illustrated in **Fig. 2a**; *G_ST_* (·) in Equation 4 and *G_DA_* (·) in Equation 7), which ensures that only the most relevant and reliable data contribute to the model training at each step. The above steps, including the gating mechanism, are repeated iteratively until the model converges. In each training step, backpropagation (31) is employed to refine the parameters of both the feature extractor and the classifier, guided by the composite loss function described earlier (Methods and SI Appendix).

For clarity and reference, the pseudo code encapsulating the entire training process of our framework is concisely summarized in **Fig. 2c**.

At last, it should be noted that we utilized all available experimental data for domain adaptation and self-training rather than dividing it into validation and testing sets, but we should also emphasize that none of the ground-truth were used throughout the training process. Our primary concern for such design is to reflect the real-world conditions of neuroscience research, where the goal is to infer connectivity from all available data without relying on ground-truth labels. Nevertheless, we have also conducted cross-animal validation experiments to evaluate our model’s generalization capabilities (see Discussions for details).

### Visualization of feature space evolution during training

Next, to gain insights into the underlying mechanisms of our DeepDAM framework, we visualized the evolution of the feature space during the training process, comparing to the conventional deep learning approach lacking domain adaptation and self-training (i.e. *vanilla* deep learning method). The synthetic dataset was generated by the MAT neural network and the experimental data were a subset of extracellular recordings from the CA1 region of freely-running mice with ground-truth labels (25) (**Fig. 3a**; SI Appendix). After training, both methods were capable of accurately distinguishing between synthetic connection and non-connection data (**Fig. 3b** bottom right) but only DeepDAM can successfully classify experimental data.

**Fig. 3.**
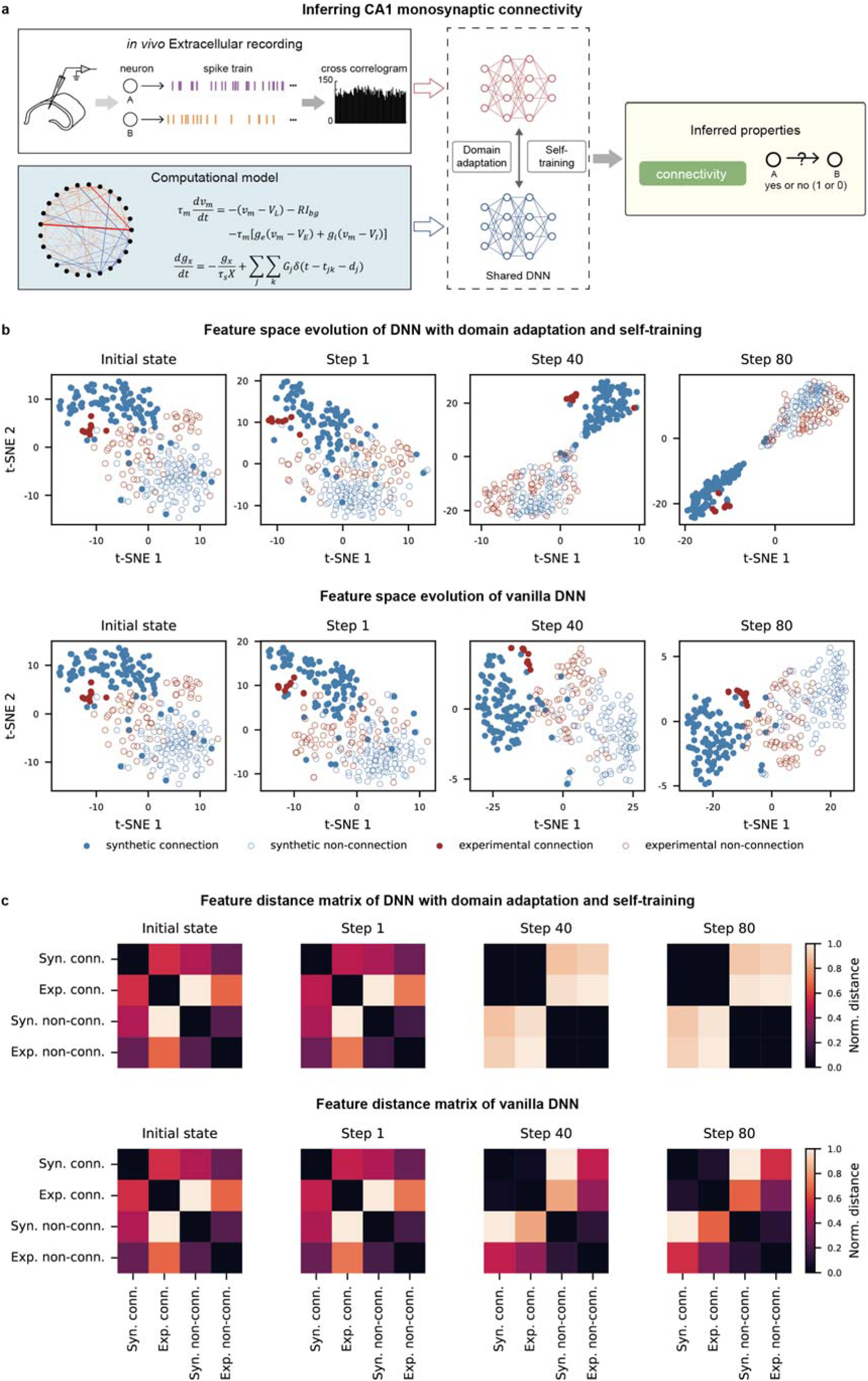
Training workflow of our inference framework. (**a**) The experimental data and computational model for inferring the existence of monosynaptic connectivity between pairs of neurons. (**b**) Evolution of feature representations across the training process for both our inference method and the vanilla deep learning method without self-training and domain adaptation. The same fifty randomly selected synthetic connection and non-connection samples are shown in each subplot. (**c**) The evolution of distance matrix of synthetic and experimental features across the training process for both our inference method and the vanilla deep learning method without self-training and domain adaptation. Distance between two groups was calculated as the average Euclidean distance between each pair of features.

For the vanilla deep learning method, the experimental data were inappropriately positioned between the synthetic data labeled as ‘connection’ and ‘non-connection’ in the feature space (**Fig. 3b** bottom right). This misplacement was due to the persistent distributional discrepancy between synthetic and experimental data in the input space, a consequence of model mismatch. As a result, the classifier tended to overfit to the synthetic data, leading to inaccurate classification of the experimental features.

In stark contrast, our method effectively utilized domain adaptation to cluster synthetic and experimental data of the same class together in the feature space (**Fig. 3b** top right). This strategic grouping allowed the classifier, which was trained on synthetic data via supervised learning and on experimental data via self-training, to accurately classify both datasets. As the training proceeds, the pseudo labels for self-training and overall prediction performance gradually stabilized (**Fig. S3**). To quantitatively evaluate the discrepancy between synthetic and experimental feature distributions, we computed the distance between these distributions (**Fig. 3c**). The results revealed that our approach successfully differentiated connection and non-connection data in the feature space, enhancing the classifier’s ability to distinguish them accurately (**Fig. 3c** top). On the contrary, the vanilla deep learning method struggled to differentiate between synthetic connection data and experimental non-connection data, and between experimental non-connection and connection data, due to model mismatch and the classifier’s overfitting to synthetic data (**Fig. 3c** bottom).

To conclude, our DeepDAM framework offers an effective solution to the model mismatch challenge, ensuring the transfer of precise neural connectivity inference from synthetic data to experimental data.

### Evaluation of DeepDAM framework’s performance on *in vivo* dataset

In this section, we systematically evaluated the performance of our framework on real *in vivo* spike data with ground-truth labels, comparing with existing popular methods. Specifically, we benchmarked against three methods: the vanilla deep learning approach, CoNNECT (12), and GLMCC (10).

Briefly, CoNNECT is essentially a deep learning method without domain adaptation and self-training, similar to the vanilla deep learning method but with a normal convolution neural network architecture. The authors of CoNNECT provided a DNN model trained solely on synthetic data generated from the same MAT network; we directly utilized their published model for performance comparison (12). GLMCC, in contrast, employed a Generalized Linear Model to fit CCGs based on synthetic data, which was then applied to rat hippocampal data. In our evaluation, we adopted their model with the original hyperparameters (10), as these were specifically tuned for hippocampal data, ensuring a fair comparison (SI Appendix).

It is essential to highlight the fundamental challenge in neuroscience research: the scarcity of *in vivo* datasets with reliable ground-truth connectivity labels. Direct measurement of synaptic connections in living brains is extraordinarily difficult with current technologies, making ground-truth connectivity largely unknown in most cases. This unique *in vivo* dataset is, to our knowledge, one of the few available with “inferred labels” provided by the dataset creators (25).

Specifically, the dataset’s labels were established using two distinct labeling methods, resulting in two subsets: (1) a smaller subset with reliable labels, validated by the consistency between the two labeling methods. This subset allows us to rigorously test the efficacy of our method on real *in vivo* data, providing a dependable benchmark for our techniques; (2) a larger subset with less reliable labels, characterized by inconsistencies between the labeling methods. This subset represents a more common scenario in real-world research, where data ambiguity is prevalent.

Our performance evaluation was based on the smaller subset with reliable labels (25). It is important to emphasize that we did not use any ground-truth labels for training our model; these labels were solely employed to assess the model’s prediction accuracy on the experimental data.

We chose MCC as our performance metric, as it is consistent with previous literature (10, 12) and also suitable for imbalanced classification problems (32). The results showed that these three existing methods exhibited noticeable limitations. Predominantly, the vanilla deep learning method tended towards false positives, while both GLMCC and CoNNECT generated a mix of false positives and negatives (**Fig. 4**). In contrast, our method overcame these limitations, achieving an MCC of 0.91 (SD=0.08), in stark contrast to vanilla deep learning (0.61, SD=0.15), GLMCC (0.62), and CoNNECT (0.58).

**Fig. 4.**
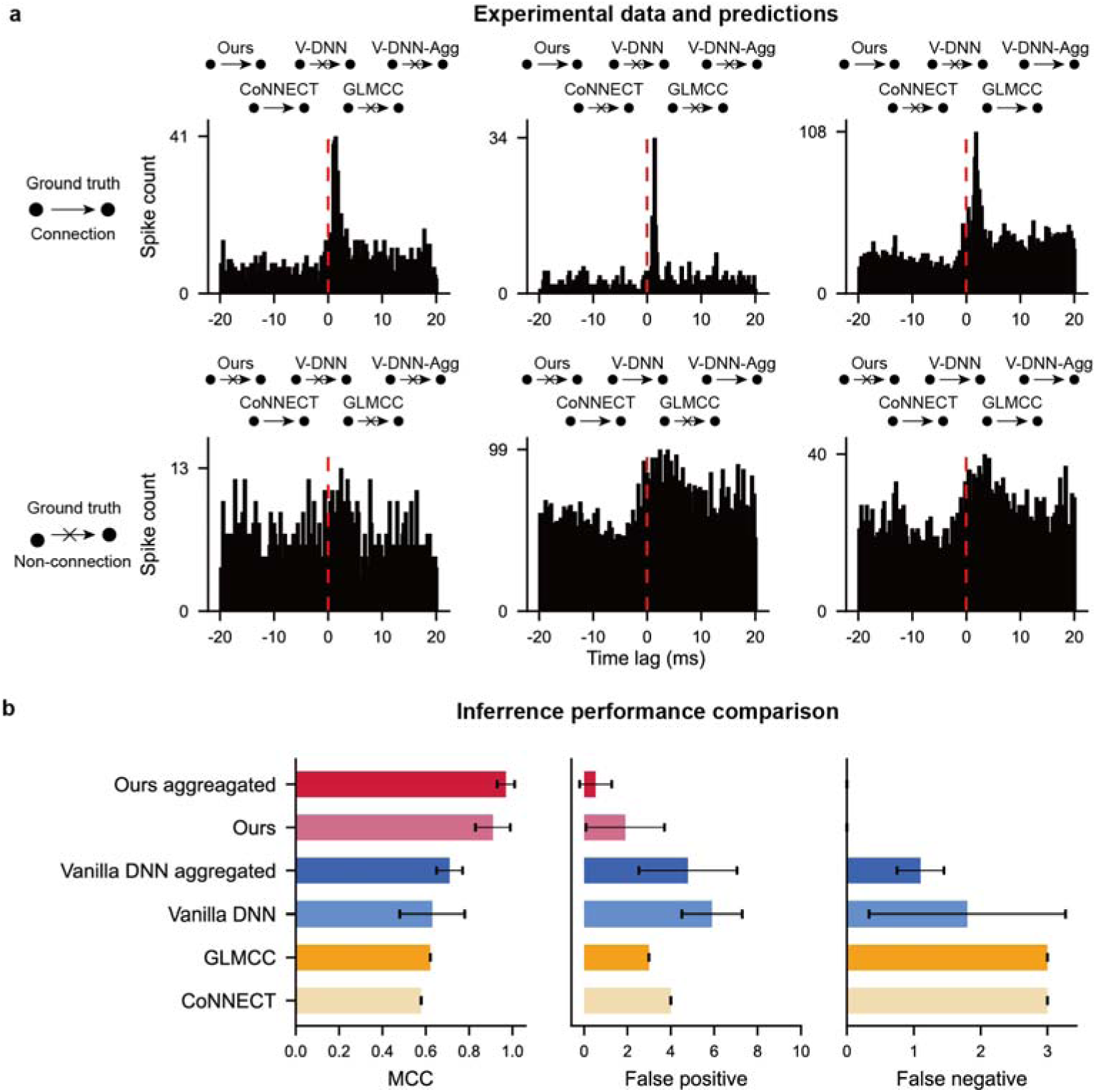
Performance comparison on experimental dataset with ground-truth labels. (**a**) Example experimental CCGs and predictions by different methods. Ours: our aggregated version. V-DNN: the vanilla deep learning method. V-DNN-Agg: the aggregated vanilla deep learning method. CCGs were plotted from [-20, 20] ms based on the neuron spike timing with the time bin of 0.2 ms. (**b**) Performance comparison between 6 different methods. Error bar: standard deviation across 10 random seeds.

Interestingly, we observed some performance fluctuations across different random seeds during the training process (**Fig. 4b**). To address this variability and ensure the robustness of our method, we introduced an aggregated version of our method, where we trained the same framework across various random initializations, generating a unique inference prediction for each experimental data point with each random seed. Subsequently, we employed a majority vote algorithm to aggregate the results from all random seeds. Remarkably, our aggregated method successfully eliminated noisy inference results and achieved an outstanding MCC of 0.97 (SD=0.04; **Fig. 4b**; Methods). Our aggregated method is fairly robust which achieved an MCC of 0.91-0.98 across a wide range of hyperparameters (**Error! Reference source not found.**) and such high performance can only be achieved with both self-training and domain adaptation (**Fig. S6**). Similarly, the hyperparameter combination we selected was also based on the consistency across random seeds (SI Appendix).

### Evaluation of DeepDAM framework’s robustness on *in vivo* dataset

A hallmark of a robust inference algorithm is its consistent performance across varied experimental scenarios. For example, the length of experimental recordings has been shown to significantly influence inference accuracy (10). Therefore, we assessed our method’s resilience by subsampling experimental recordings to varying lengths. As recording length increased, our method’s performance consistently improved (**Fig. 5b**). This is likely due to longer recordings capturing more nuanced neuron interactions, whereas shorter lengths can obscure critical temporal patterns (**Fig. 5a**). Impressively, even when restricted to 20-minute recordings, our method surpassed other frameworks tested on full-length recordings (0.15-0.27 higher MCC score) (**Fig. 4b** & **Fig. 5b**).

**Fig. 5.**
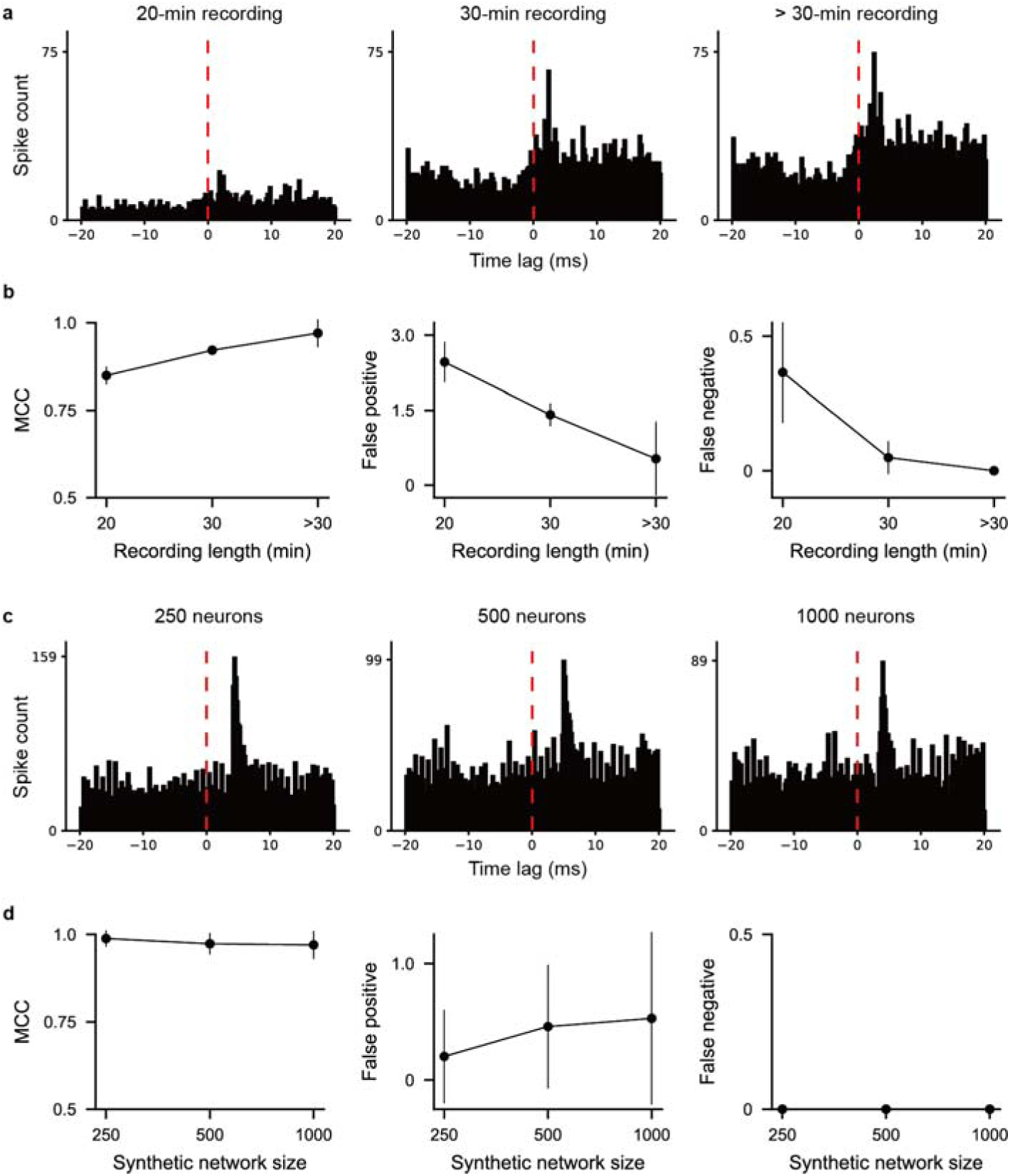
Performance stability across different experimental conditions and computational model structures. (**a**) Experimental CCG of the same pair of neurons at different recording lengths. (**b**) MCC, false positives, false negatives of our aggregated method at different recording lengths. Error bar: standard deviation across 10 random seeds. (**c**) Synthetic CCGs generated from MAT neural networks of different sizes. (**d**) MCC, false positives, false negatives of our aggregated method at different sizes of MAT neural networks. Error bar: standard deviation across 10 random seeds.

Furthermore, considering the variability in computational models employed in different studies, we tested our method’s robustness by training it using three MAT neural networks of different sizes (*N_neuron_* = 250, 500, 1000, respectively). The consistent outcome was a robust MCC of 0.99 for *N_neuron_* = 250 (SD=0.02), 0.97 for *N_neuron_* = 500 (SD=0.03), 0.97 for *N_neuron_* = 1000 (SD=0.03), across all models (**Fig. 5d**), highlighting our framework’s adaptability across diverse research scenarios. Interestingly, the highest performance, though marginally, was achieved at *N_neuron_* = 250, suggesting that this size of network model might best mimic the dynamics of the recorded neural ensemble.

We conducted a detailed analysis of the failure cases of all tested methods. Notably, our techniques only resulted in false positives, which were potentially caused by three different reasons. The first reason is the covariation between pairs of neurons (blue examples in **Fig. S4**), where the center peak starts before 0s and the amplitude of the peak is not significantly larger than the baseline fluctuation. Secondly, insufficient number of spikes can lead to false negatives (purple examples in **Fig. S4**). The third reason is that the CCGs for these cases exhibited small peaks just after zero lag (orange examples in **Fig. S4**), which likely led DeepDAM to misclassify them as connections. These concerned CCGs displayed slight peaks after zero, likely causing DeepDAM to incorrectly classify them (**Fig. S4**). Furthermore, these CCGs generally had fewer spikes compared to the clearer CCGs in **Fig. 4**, supporting our observation that longer recording durations tend to enhance performance. We highlighted that the aggregated DeepDAM only failed four CCGs in all 100 aggregations, demonstrating the robustness of our approach. Conversely, other methods displayed a mix of false positives and false negatives. For the false positives of all other methods, some of them are also due to the lack of spikes (purple examples in **Fig. S4**). While for other mistakes, interestingly, they are largely affected by covariations between pairs of neurons, which widely exist in the neural system, such neural oscillations and common inputs. Strikingly, all other methods made false negatives even if there is a very clear peak after 0s in the CCGs (right two columns from 3^rd^ to 6^th^ rows of **Fig. S4**). This suggests that methods developed solely on synthetic data have limited generalization capabilities on real experimental data.

To conclude, our DeepDAM framework, which synergizes domain adaptation and self-training, effectively mitigates the model mismatch problem and delivers outstanding performance in inferring monosynaptic connectivity from *in vivo* spike trains. Its demonstrated robustness, evident in handling varying recording lengths and accommodating different model structures, establishes the framework as a versatile and powerful tool, offering significant potential for broader applications in the field.

### Application to a broader *in vivo* spike dataset with low signal-to-noise ratios

As we transition our method to real-world scenarios, its capability to effectively handle uncertain or noisy data becomes increasingly important. In the previous section, our framework was tested on the subset from English et al. (25), where ground-truth labels were consistently established by two labeling methods, suggesting that the CCGs from this subset typically exhibit high signal-to-noise ratios. To further evaluate the robustness of our framework, we expanded our analysis to include a broader set of *in vivo* spike data characterized by lower signal-to-noise ratios. This larger dataset, also sourced from English et al.’s study (25), contained CCGs with inconsistent or uncertain ground-truth labels according to the two tagging methods, which is a scenario likely more common in *in vivo* experiments (**Fig. 6a**; SI Appendix).

**Fig. 6.**
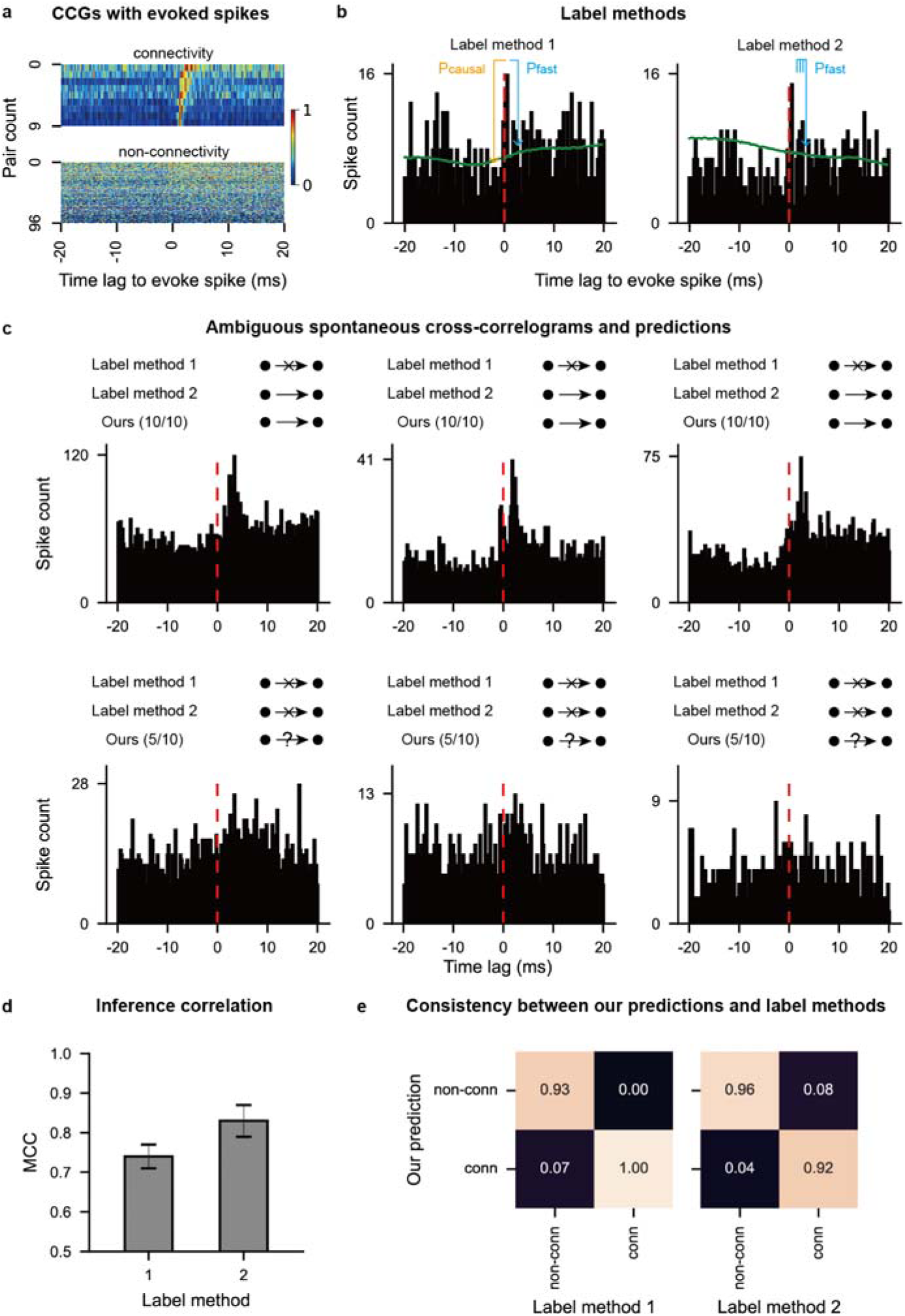
Consistency between the prediction of our aggregated method and two label methods. (**a**) All CCGs with evoked spikes in the large dataset. For labeling, the CCGs were calculated based on ‘presynaptic’ spikes that were evoked via juxtacellular stimulation. Spontaneous spikes of the ‘presynaptic’ neuron were discarded. The label is given by label method 1. Color bar denotes the spike count, normalized by the maximal value of all CCGs. (**b**) Illustration of two label methods. The green line represents the slowly co-modulated baseline (SI Appendix). Briefly, label method 1 tests the lowest p-value of time bins in both directions (*p*_causal_ and *p*_fast_) while label method 2 tests the p-value of three consecutive time bins in one direction (SI Appendix). (**c**) Experimental CCGs and predictions of our aggregated method and two label methods. For inference, CCGs were calculated using spontaneous spikes of the ‘presynaptic’ neuron, while evoked spikes and the corresponding stimulus window was discarded from the CCGs. The numbers in the parentheses indicate the count of random seeds out of 10 that resulted in the same inference outcome. (**d**) MCC between our aggregated method and two label methods on the amibigous dataset. (**e**) Consistency matrix between our aggregated method and two ground-truth tagging methods on the amibigous dataset. For example, the upper-left grid in the left panel represented the percentage of experimental CCGs in the dataset that both our aggregated method and label method 1 inferred as non-connections.

Interestingly, in the enlarged experimental dataset, our aggregated method demonstrated greater consistency with the second ground-truth tagging method, which tended to identify connections more frequently than the first method (**Fig. 6b & c**; Discussions). Yet, discrepancies still existed between our method and both tagging methods, requiring further investigation to determine the most effective approach. Our previous evaluations suggest that increasing the recording time, reducing CCG noise and enhancing prediction reliability could be a viable solution (**Fig. 5b**). Additionally, we encountered cases where neither tagging method could definitively classify certain CCGs as connections or non-connections due to insignificant statistical tests. Our inference method also exhibited uncertainty in these cases (**Fig. 6a** bottom row), primarily due to these CCGs having an insufficient number of spikes for the corresponding neuron pairs (**Fig. 6a** bottom row), underscoring the need for longer recording times for neurons with lower firing rates to yield convincing inferences (**Fig. 5b**) (10).

In conclusion, our findings emphasize the need for comprehensive evaluations using large-scale datasets with accurate ground truths to refine and enhance connectivity inference methods. Improving the ground-truth labeling process and expanding the dataset size are essential steps to enhance the robustness of the inference methods.

### Versatility in inferring biophysical properties across neural scales

*In vivo* datasets that pair spike trains with ground-truth synaptic connectivity are extremely scarce, limiting our ability to directly validate connectivity inference methods. In fact, to our best knowledge, only a single *in vivo* spike recording dataset with known synaptic connections (from mouse hippocampal CA1) is available, which makes comprehensive performance evaluation challenging. To address this and demonstrate DeepDAM’s effectiveness on broad real experimental data, we turn to *ex vivo* and *in vitro* preparations as alternative validation platforms. In this final section, we showcase DeepDAM’s adaptability under broader model-mismatch conditions across different recording modalities, species, and observation scales, by applying it to traditional model-fitting tasks. This indirect evaluation strategy, using controlled laboratory datasets, supports the generalizability of our approach despite the paucity of fully-labeled *in vivo* benchmarks.

Based on the foundational Equations 1–4 of our framework, we can extend DeepDAM beyond connectivity mapping to infer detailed biophysical properties of single neurons and microcircuits. The key modification required is a subtle adjustment of the discrepancy loss function *L_dis_*, (as described SI Appendix) to target specific observable features. Unlike the model-generation paradigm used earlier for connectivity inference, where synthetic data are generated from a fixed model, here we employ a model-fitting paradigm. In these tasks, model parameters are iteratively optimized so that the model’s responses closely align with experimental observations in terms of the chosen biophysical features (SI Appendix). This shift from data generation to model fitting illustrates DeepDAM’s versatility: with minimal changes to the loss function, the same framework can capture neuron-level properties (e.g. membrane dynamics) or circuit-level behaviors from experimental recordings, further demonstrating its robustness across neural scales.

As for the single-neuron inference, our framework inferred biophysical properties of single neurons of mice using intracellular recordings *in vitro*, including passive membrane dynamics, background noise magnitude, and attributes of ion channels. Such properties align with the parameters set by the single-compartment Hodgkin-Huxley model, known for capturing the vast dynamics of cortical neurons (**Fig. S8a**; SI Appendix) (33). We assessed the z-scored mean absolute error (MAE) between biophysical patterns of synthetic model responses and experimental data, including spiking statistics and membrane potential behaviors (**Fig. S8d**). Impressively, our method efficiently inferred biophysical patterns of 574 neurons across 14 brain regions in the Allen Cell Types Dataset (22) (**Fig. S9a** & **b**), with consistent inference accuracy across different regions. The results revealed that both spike and subthreshold patterns, which were major distinguishable patterns across different neuron types, were well predicted, suggesting that our inference results could potentially facilitate studying subcellular differences among different neuron types. Among all biophysical patterns, the half-spike width is the most poorly tuned biophysical pattern (**Fig. S9c** & **d**). One potential reason is that the single-compartment HH model is relatively simple to fit this biophysical pattern, while increasing the number of (sodium) channels could give a better fitting accuracy (21). Moreover, when compared with evolutionary search (ES) algorithm, perhaps the most popular method for single-neuron biophysics inference (23), our approach demonstrated superior performance, achieving an inference score 1.89 times higher (i.e., number of good inferences given the same sample size; our framework: 574, ES: 303, total: 1052) (**Fig. S9e**; SI Appendix).

Furthermore, our framework inferred 31 distinct properties of the intricate stomatogastric ganglion microcircuit of the *Cancer Borealis* (**Fig. S10a**), including nuances of enriched ion channels and diverse synapses, which aims to replicate the circuit’s characteristic pyloric patterns (**Fig. S10b**, **c** & **d**; SI Appendix). To compare the inference performance, we evaluated it against the state-of-the-art sequential neural posterior estimation (SNPE) method, specifically SNPE-C (34) (performance given by the pretrained model from Gonçalves et al. (23); SI Appendix). Our framework demonstrated a 2.54 times smaller inference error for all biophysical patterns than SNPE-C (MAE of our framework: 0.0136, SD=0.0024; MAE of SNPE-C: 0.0346, SD=0.0048) (**Fig. S10e**). In summary, the above results underscore our framework’s robustness and adaptability in neural analysis across multiple scales.

## Discussion

The present study introduced a novel model-based deep learning inference framework to tackle the challenging problem of neural circuit connectivity reconstruction from *in vivo* recordings, which could potentially bridge the gap between neural circuitry functions and its architecture (1-4, 35, 36). It has been demonstrated that our framework overcame the limitations of existing model-based methods (10, 12) and accurately inferred monosynaptic connectivity in the hippocampal CA1 region of freely running mice. The key innovation of our approach lies in its ability to address the challenges posed by the model mismatch problem, which often leads to biased inference outcomes (8). By integrating domain adaptation and self-training with computational models, our framework adapts to discrepancies between synthetic and experimental data and establishes a mapping from paired spike trains to their corresponding connectivity, significantly improving the accuracy of connectivity inference. We have also demonstrated the broad applicability of our framework in inferring biophysical properties of single neurons and microcircuits. By extending the framework to classic model fitting tasks, we successfully inferred a range of biophysical properties of single neurons in the mouse primary visual cortex and the stomatogastric ganglion microcircuit of the *Cancer Borealis*. In both cases, our approach outperformed existing methods in accuracy and efficiency (21, 23), highlighting its potential for advancing the study of neural computation and function across various observational scales.

One of the major challenges in inferring neural connectivity from *in vivo* recordings is the partial observation of neurons, leading to incomplete knowledge of the network. As a result, computational models for the biological network often fail to capture the repertoire of the real biological dynamics and could produce substantial model artefacts. Eliminating such mismatch based on prior knowledge is difficult due to the intricate complexity and insufficient understanding of the underlying biological mechanisms. Consequently, this unrecognized mismatch can introduce bias in connectivity inference when solely relying on the model as the reference (8). Our framework effectively addressed these obstacles by learning domain-invariant and informative features through domain adaptation and fine-tuning the connectivity classifier using experimental data through self-training. As a result, our method achieved 0.97-1.0 MCC to infer monosynaptic connectivity of the mice CA1 region on a dataset with high signal-to-noise ratio, significantly exceeding current methods. Notably, these existing methods all demonstrated near-perfect inference results on synthetic data (10, 12) but failed to transfer such excellent performance to real experimental data. Similar trend was seen in model-based inference framework in other domains, such as inferring biophysical properties of single neurons and microcircuits (13, 37). Our study not only formulated this issue as the OOD problem in machine learning and proposed a unified solution, but also emphasized the importance of evaluating model-based inference techniques using real experimental data.

Another straightforward and widely used method for tackling the mismatch problem in robotics and autonomous driving is domain randomization (38–40). This technique involves varying model parameters to enhance the diversity of synthetic data from different domains. This could potentially enable the machine learning model to learn domain-invariant features that can generalize to real-world applications. Therefore, to test whether increasing the diversity and size of the synthetic dataset can lead to better inference, we simulated 27 different MAT networks by altering three keys: background noise, connectivity ratio, and network size to generate a synthetic dataset that was 16.5 times larger than the previous one (SI Appendix). Our tests revealed that domain randomization increased the MCC by 0.11 for the vanilla deep learning method and by 0.03 for its aggregated version. Despite these improvements, the performance still lagged behind that of DeepDAM and its aggregated version by 0.17 and 0.23 MCC respectively (**Fig. S5**). These results suggest that while domain randomization can enhance the performance of standard deep learning approaches, the benefits are relatively modest, likely due to the high-dimensional nature of the model and the complexity of the neural structure involved. Interestingly, domain randomization can also improve the performance of DeepDAM to an MCC score of 1.0, suggesting that combining DeepDAM with other advanced machine learning techniques is a promising future direction.

The development of large-scale datasets with ground-truth connectivity is urgently needed to further assess and compare these methods, especially because different methods yield varying inference predictions on ambiguous data (**Fig. 6**) and the performance of our method has already saturated on this available dataset. For these ambiguous data, our aggregated method was generally more consistent with the second modified label tagging method (**Fig. 6d**). We introduced this modified label tagging method because we observed that the experimentally validated ground-truth tagging method was excessively strict. It demanded the peak of the CCG to significantly surpass the slowly co-modulated baseline by a considerable margin (*p*_fast_ threshold: 0.001) and to be significantly higher than the peak in the anti-causal direction (**Fig. 6b**) (25). We relaxed the *p*_fast_ threshold to 0.01 for three consecutive time bins and eliminated the requirement in the anti-causal direction (i.e., no *p*_causal_ prerequisites; **Fig. 6b**; SI Appendix), so the adjusted tagging method leaned toward identifying more connections. The fact that our aggregated inference method aligned closer with the modified tagging method (**Fig. 6c** top row) suggests that the original labeling criterion often led to the classification of numerous CCGs with relevant connectivity information as indeterminate, such as bidirectional connections or those coexisting with network modulations (**Fig. 6c** top row). Mitigating the impact of these ambiguous CCGs can be accomplished in several ways. Firstly, extending the recording duration (**Fig. 5b**) can be a straightforward approach, as many experimental methods support chronic recordings of the same group of neurons (7). Alternatively, incorporating probabilistic methods (41), ensemble methods (42) or noisy pseudo labels (16, 18, 19) into our existing framework can explicitly model the uncertainty in our inference results of such ambiguous cases.

In this study, we tailored the DeepDAM training process to better address real-world experimental requirements. We included all experimental data for domain adaptation and self-training without using any ground-truth labels during training. We chose not to segregate data from different animals for evaluation primarily for two reasons: 1) Ideally, in real-world settings, we aim to analyze all available data rather than just a subset; 2) Variations in data across subjects can reduce inference accuracy, similar to the mismatch observed between synthetic and experimental data. Addressing the issue of cross-subject generalization, which remains a challenge in the field, we assessed the generalization ability of our DeepDAM framework using a “leave-one-animal-out” approach across 8 mice. The generalization performance of both DeepDAM (MCC: 0.88, SD=0.15) and its aggregated version (MCC: 0.95, SD=0.07) was remarkably close to their initial performance (DeepDAM MCC: 0.91, SD=0.08; DeepDAM aggregated MCC: 0.97, SD=0.03) and substantially better than other methods, including those employing domain randomization (**Fig. S7**). These results underscore the adaptability and effectiveness of DeepDAM in handling the inherent variability found in real-world experimental settings.

We evaluated DeepDAM’s performance in recovering monosynaptic connections within hippocampal CA1, but its applicability to other areas remains to be determined. Because neural circuits differ fundamentally across regions—giving rise to unique physiological processes and activity signatures (43)—DeepDAM relies on a computational model that captures those region-specific firing dynamics, and our use of the MAT network is tailored to CA1 properties. Going forward, extending DeepDAM will require generating ground-truth spiking datasets from diverse brain regions and developing biophysical models that faithfully reproduce each region’s electrophysiological characteristics.

Our results also demonstrated the robustness and general applicability of the DeepDAM framework, which is designed to be agnostic to other diversity of the neural system. The framework depends on a mechanistic model capable of approximating the dynamics of the underlying systems, such as the firing rate distribution. Therefore, once such simple mechanistic model has been obtained in another species, such as the visual cortex in primates (44), the DeepDAM framework can be easily applied. While it stands to gain from advancements in more precise mechanistic models, its potential significance lies in advancing the mechanistic modeling of large-scale neural circuits. This is crucial because commonly used techniques like artificial recurrent neural networks (45) and multicompartmental neuronal networks (46, 47) often encounter significant model mismatch issues, which can also be conceptualized as an OOD problems (48). Therefore, we anticipate that DeepDAM, in conjunction with these modeling techniques, will promote a closed-loop, iterative process for designing and refining mechanistic models to directly deduce the system’s architecture and biophysics from experimental data.

Furthermore, DeepDAM can be easily extended to the classic model-fitting task for biophysical inference of single neurons and microcircuits through changing the discrepancy loss and achieved high accuracy and efficiency in both cases. One limitation of DeepDAM on model fitting is that it can only output one set of feasible neural properties to match the experimental data, while many studies have demonstrated there can be multiple sets of feasible properties (23, 49-54). This could be potentially achieved by replacing the *L_syn_* with the loss function used in SNPE methods (23).

In summary, our framework substantially alleviates the OOD problem in inferring monosynaptic connectivity and biophysical properties from experimental data. This indicates that our framework can potentially facilitate the interaction between experimental and modeling studies to uncover the relationship between low-level neural properties and high-level functions. With continued advancements in experimental techniques and machine learning methodologies, our framework paves the way for a more profound understanding of the brain’s intricate mechanisms, leading to transformative insights into neural computation and cognition. For example, our framework explores the mechanistic formation of neural representations underlying behavior and cognition from a bottom-up perspective, providing a valuable complement to current statistical and geometrical analysis on large-scale neural representations from a top-down perspective (55–57). Furthermore, our framework is in line with the rising next-generation ML techniques that utilize simulators and synthetic data to learn richer representations of real world (39). By tightly fusing computational modeling and ML techniques, our framework may inspire ground-breaking methods that use enriched computational models and synthetic data to address cutting-edge challenges in multiple fields (39, 58), such as behavioral analysis (59), protein function prediction (60), and medicine and healthcare (61).

## Methods

Here we describe the training details of the DeepDAM framework and leaves other methodological details in SI Appendix.

### Training configurations to infer monosynaptic connectivity

As our method is purely unsupervised and does not use any ground-truth labels, the entire experimental dataset was used for domain adaptation and self-training during the training process. The batch size is 800. The initial learning rate is 0.0001 and adapted by the Adam optimizer (62) with *β*_1_ = 0.9 and *β*_2_ = 0.999. The discrepancy loss is defined as the estimated contrastive domain discrepancy at the feature space (14), given by Equation 5-7:

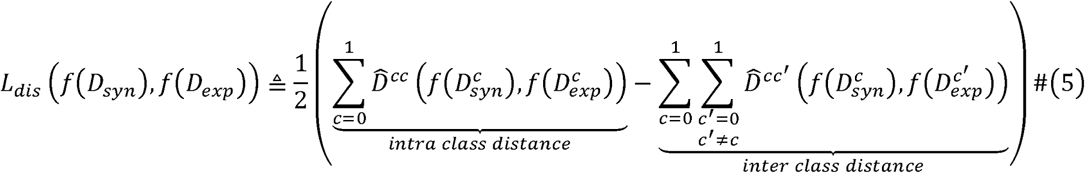

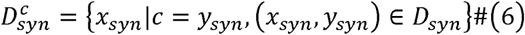

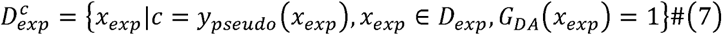

where *G_DA_* (·) is the binary gating function for domain adaptation and 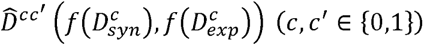 is the empirical kernel mean embedding which describes the distributional distance at the feature space between synthetic data labeled as class *c* and experimental data pseudo-labeled as class *c’* The empirical kernel mean embeddings of the contrastive domain discrepancy are given by Equation 8-12:

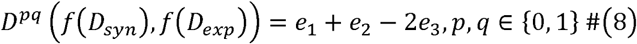

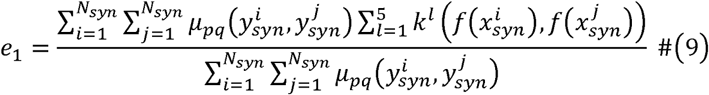

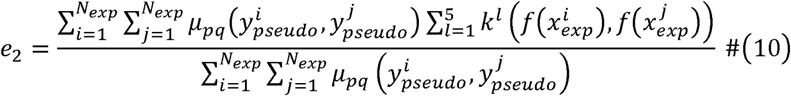

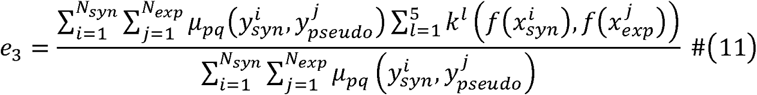

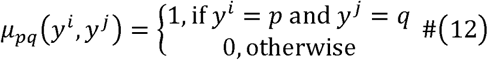

where *k^l^* is a Gaussian kernel with a variance of 2*^l^*normalized by the number of samples and total distance (14). The discrepancy scaling factor is set as 0.001. The gating threshold for eliminating experimental data from domain adaptation is 0.01 and the gating threshold for self-training is 0.01.

Pseudo labels of experimental data are obtained by applying the spherical K-means clustering algorithm at the feature space among synthetic data and experimental data. The initial cluster center of class is the synthetic cluster center, given by Equation 13. Then the algorithm repeats the following two steps until convergence or reaching the maximum iterations: 1) assigning each experimental datum a pseudo label based on the nearest cluster center; 2) updating the cluster center based on experimental data, given by Equation 14.

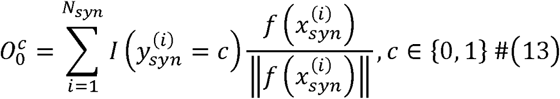

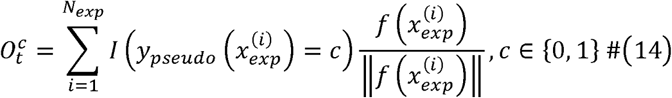

where 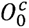 is the initial cluster center for class 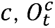 is the cluster center for class *c* at training step *t* and *I* is the indicator function. Crucially, synthetic data labels ensure that synthetic data with different labels are not grouped in the same cluster.

As the nature of the pseudo labels is noisy, we incorporated the generalized cross-entropy loss (Equation 15) which has been demonstrated to be noise-robust (20).

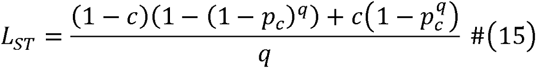

Where *p_c_* = *g*(*f*(*x_exp_*)) is the output probability for class*c* of DeepDAM. In this study, we chose *q* = 1

For the aggregated method, we trained the DeepDAM framework for 10 different random seeds. The we ensembled the predictions from 5 randomly selected seeds and used the mean probability for each class as the aggregated probability to compute our performance metrics. This procedure was repeated 100 times.

## Author contributions

Conceptualization and methodology: KS, KD

Investigation and analysis: KS, SZ, MB, PQ, LY

Discussion: KS, SZ, MB, PQ, LY, XL, LH, YZ, LM, KD

Supervision: KD, LM

Writing—original draft: KS, SZ, MB, LM, KD

## Data and materials availability

The source code of our framework in this work will be available upon acceptance. All data are available in the main text or the supplementary materials.

