## Supplementary material for "Domain-adaptive matching bridges synthetic and *in vivo* neural dynamics for neural circuit connectivity inference": SI Appendix

Supporting Information

**Deep neural network architecture**

The chosen architecture for the temporal feature extractor is the ResNet-LSTM, renowned for its prowess in processing high-dimensional temporal data (1). The temporal feature extractor first extracts local temporal features by ResNet and then extracts global temporal dependencies of the local features using LSTM cells (**Fig. 2**). Residual Blocks (ResBlocks) are the basic components in ResNet-50 and other variants of ResNet. Each ResBlock comprises three units, each containing a convolutional layer (whose kernel size is 1, 3, 1 correspondingly), a batch normalization layer, and a leaky ReLU activation layer. A skip connection associates the input with the output of the ResBlock through a summation operation. Compared with the original ResNet-50 design, we modified the first input channel from 3 to the number of model response sequences, replaced the ReLU activation function with the Leaky ReLU activation function, and replaced the Batch Normalization layers with the Group Normalization layers with 32 groups (2), as Batch Normalization was unstable for data from a different statistical domain (3). We also added another convolutional layer with kernel size of 1 and dimension of 512 before the final average pooling layer. The number of hidden units of LSTM is 2048. The classifier consists of 1 fully connected layer.

**Hyperparameter selection**

Given the performance fluctuations across random seeds, we selected the hyperparameter setting based on the consistency across different random seeds for aggregated DeepDAM. Specifically, for each aggregation, we obtained the predicted connectivity for all pairs of experimental CCGs. Then we calculated the mean Pearson correlation coefficient between the predicted connectivity vector of each aggregation. The hyperparameter which gives the highest correlation, indicating the most consistent prediction across random seeds, was selected as the final setting.

**Comparison with CoNNECT and GLMCC on the inference of neural connectivity**

GLMCC method is proposed by Kobayashi et al. (4) and we used the published code at <https://github.com/NII-Kobayashi>, where hyperparameters remain unchanged as in the original study (4). For example, we set the membrane time constant as 5 ms, and $\gamma$ as $5\times{10}^{-4}$, which are identical to the original study. The time length for a cross-correlogram is $100$ ms and the time bin is $1$ ms. Synaptic delay is tested from 1-5 ms at the step of 1 ms and selected as the one that gives the highest likelihood. As for CoNNECT, we used the pre-trained model obtained from [https://github.com/shigerushinomoto/CoNNECT](https://github.com/shigerushinomoto/CoNNECT.git) and set the classification threshold as 0.5 (5).

**Ablation analysis**

To evaluate the performance of the two key components in the DeepDAM framework: domain adaptation and self-training, we performed ablation studies on the two components. The highest performance of each ablated version was selected for comparison with our DeepDAM framework.

**Domain randomization for DeepDAM and vanilla deep learning methods**

We conducted a parameter sweep across three hyperparameters of the MAT network, involving the proportion of neurons receiving background inputs at levels of 0.05, 0.1, and 0.25, the connectivity rate for each neuron linking to excitatory neurons at 0.0625, 0.125, and 0.25, and network sizes of 250, 500, and 1000. This created 27 distinct networks, generating a total of 6,361,076 pairs of CCGs. We then trained both our DeepDAM framework and vanilla deep learning models using this expanded synthetic dataset under the same conditions as previously used.

**Dimension reduction and visualization with t-SNE**

For visualization with t-SNE in **Fig. 1** and **Fig. 3**, the dimension of original data or the high-dimensional features was first reduced by principal component analysis to 50. The distance metric for t-SNE is Euclidean distance and the perplexity of t-SNE is set as 30. In each processing, the randomly selected 50 synthetic connection data and 50 synthetic non-connection data were the same.

**Large-scale MAT neural network model**

We used the MAT neural network as the computational model, which is more biologically realistic than the Hodgkin-Huxley neural network in terms of fluctuated inputs (5). The membrane potential of a neuron is driven by synaptic inputs and background noises, given by Equation S1. The synaptic conductance evolves with time, given by Equation S2. Excitatory synaptic weights are drawn from a log-normal distribution, given by Equation S3. Inhibitory synaptic weights are drawn from a normal distribution, given by Equation S4. Only excitatory synaptic connections with $G\geq0.01$ are included in the training set as weak connections cannot be inferred when the recording/simulation time is not long enough (4). When scaled to different network sizes, the ratio between excitatory and inhibitory neurons is set fixed to be 4:1.

$$\begin{aligned} \tau_{m}\frac{dv_{m}}{dt}=-\left( v_{m}-V_{L} \right)-\tau_{m}\left[ g_{e}\left( v_{m}-V_{E} \right)+g_{i}\left( v_{m}-V_{I} \right) \right]-RI_{bg}\#(S1) \end{aligned}$$

$$\begin{aligned} \frac{dg_{x}}{dx}=-\frac{g_{x}}{\tau_{s, x}}+\sum_{j} \sum_{k} G_{j}\delta\left( t-t_{jk}-d_{j} \right)\#(S2) \end{aligned}$$

$$\begin{aligned} P\left( G_{exc} \right)=\frac{1}{\sqrt{2\pi}\sigma_{inh}}\exp\left( -\frac{\left( \log G_{exc}-\mu_{exc} \right)^{2}}{2\sigma_{exc}^{2}} \right)\#(S4) \end{aligned}$$

$$\begin{aligned} P\left( G_{inh} \right)=\frac{1}{\sqrt{2\pi}\sigma_{inh}}\exp\left( -\frac{\left( G_{inh}-\mu_{inh} \right)^{2}}{2\sigma_{inh}^{2}} \right)\#(S4) \end{aligned}$$

where $\tau_{m}$ is the membrane time constant, $v_{m}$ is the membrane potential, $V_{L}$ is the leaky reversal potential, $g_{e}$ and $g_{i}$ are excitatory and inhibitory conductances, $V_{E}$ and $V_{I}$ are excitatory and inhibitory synaptic reversal potentials, $RI_{bg}$ is the background noise, $\tau_{s, x}$ is the excitatory or inhibitory synaptic time constant, $\delta$ is the Dirac delta function, $t_{jk}$ is the $k$th spike of the $j$th neuron, $d_{j}$ is the synaptic delay, and $G_{j}$ is the synaptic weight. For full model details and parameter settings, see Endo et al. (5).

**Ground truth of monosynaptic connections *in vivo***

The experimental data are juxtacellular-extracellular recordings from the hippocampal CA1 region in awake mice (6). We only considered excitatory synaptic connections. Neurons whose firing rates are less than 3 Hz and spikes during ripple events are eliminated from the analysis. In order to mimic normal experimental conditions, all CCGs in both datasets are calculated from spontaneous spikes, meaning that all juxta-cellularly evoked spikes of pyramidal neurons are eliminated. The raw experimental data can be found at [https://buzsakilab.nyumc.org/datasets/McKenzieS](https://buzsakilab.nyumc.org/datasets/McKenzieS/).

For the first ground-truth tagging method, the time bin for CCGs is $0.4$ ms and the length was $200$ ms. Two criteria are considered for determining the presence of synaptic connections (6). The peak of the CCG in the interval $[0.8, 2.8]$ ms should firstly be compared to the slowly co-modulated baseline (Equation S5), and then compared to the largest peak in the anti-causal direction (Equation S6). The lower frequency baseline $\lambda_{slow}$ is the CCG convolved with a partially hollow Gaussian kernel, with a standard deviation of $10$ ms and a hollow fraction of 60%. The probability to get an observed (or higher) synchrony count in the $m^{\mathrm{th}}$ time lag of the cross-correlogram given $\lambda_{slow}$ in the same bin is estimated using Poisson distribution with a continuity correction.

Pairs that are considered as synaptic connections should have $P_{fast}<0.001$ and $P_{causal}<0.0026$, and that are considered as absent connections should have $P_{fast}>0.15$ and $P_{causal}>0.15$ for the small dataset used for comparison of inference accuracy (**Fig. 4** & **Fig. 5**), while $P_{fast}>0.1$ and $P_{causal}>0.1$ for the larger dataset used in **Fig. 6**. The script for generating the labels can be found at <https://github.com/buzsakilab/buzcode>.

$$\begin{aligned} P_{fast}\left( n or more | \lambda_{slow}\left( m \right) \right)=1-\sum_{x=0}^{n-1} \frac{e^{-\lambda_{slow}\left( m \right)}\lambda_{slow}\left( m \right)^{x}}{x!}-\frac{e^{-\lambda_{slow}\left( m \right)}\lambda_{slow}\left( m \right)^{n}}{2n!} \#(S5) \end{aligned}$$

$$\begin{aligned} P_{causal}\left( n or more | \lambda_{anticausal}\left( -m \right) \right) \\ =1-\sum_{x=0}^{n-1} \frac{e^{-\lambda_{anticausal}\left( -m \right)}\lambda_{anticausal}\left( -m \right)^{x}}{x!}-\frac{e^{-\lambda_{anticausal}\left( -m \right)}\lambda_{anticausal}\left( -m \right)^{n}}{2n!}\#(S6) \end{aligned}$$

This first ground-truth tagging approach is quite conservative and can miss some valid CCGs. Typically, it filters out CCGs with a monosynaptic connection profile ($[0.5, 2]$ ms peak) superimposed over a common input profile ($0$ ms peak) because it does not allow any significant modulation in the anti-causal direction (between $-2$ and $0$ ms). Therefore, we complement this approach with a second more inclusive test. As above, the peak of the CCG in the interval $[0.8, 2.8]$ ms should be significantly different from the slowly co-modulated baseline $\lambda_{slow}$ ($5$ ms wide hollow Gaussian kernel) but using a $P_{fast}$ threshold of 0.01 rather than 0.001 for 3 consecutive $0.3$ ms bins, and without any consideration for time bins in the anti-causal direction.

We constructed two datasets of experimental CCGs. For the first small dataset used for comparison of inference accuracy, the two ground-truth tagging methods give out the same labels for all CCGs. There are 9 connection data and 66 no-connection data. For the second large dataset, there are discrepancies between the results of the two methods. There are 105 data in total. The first method gives out 9 connection data and 96 no-connection data, while the second method gives out 13 connection data and 92 no-connection data.

**HH single-compartment neuron model**

The HH single-compartment neuron model is given by Equation S7-S8.

$$\begin{aligned} C_{m}\frac{dV}{dt}=g_{L}\left( E_{L}-V \right)+\bar{g}_{\text{Na}}m^{3}h\left( E_{\text{Na}}-V \right)+\bar{g}_{\text{K}}n^{4}\left( E_{K}-V \right)+\bar{g}_{M}p\left( E_{K}-V \right)+I\left( t \right)+\sigma\eta\left( t \right)\#(S7) \end{aligned}$$

$$\begin{aligned} \frac{dq}{dt}=\frac{q_{\infty}\left( V \right)-q}{\tau_{q}\left( V \right)},\quad q\in\left\{ m,h,n,p \right\}\#(S8) \end{aligned}$$

where $C_{m}$ is the membrane capacity, $E_{\text{Na}}=53$ mV, $E_{\text{K}}=-107$ mV and $E_{\text{L}}$ are reversal potentials, $V_{0}$ is the initial membrane potential, $V_{T}$ is the spike threshold, $\bar{g}_{Na}$, $\bar{g}_{K}$ and $\bar{g}_{M}$ are the maximal conductance of each ion channel, $g_{L}$ is the leaky ion channel conductance, $\tau_{\text{max}}$ is the time constant scaling factor, $\sigma$ is the noise factor, and the intrinsic noise $\eta\left( t \right)$ is the standard Gaussian noise. There are 8 biophysical properties to infer, which are $g_{\mathrm{Na}}$, $g_{\text{K}}$, $g_{\text{L}}$, $g_{\text{M}}$, $\tau_{\text{max}}$, $V_{\text{T}}$, $\sigma$and $E_{\text{L}}$. The lower bound is $[0.5, {10}^{-4} , {10}^{-4} , {10}^{-4} , 50, -90, {10}^{-4} ,-100]$ and the upper bound is $[80, 15, 0.6, 0.6, 3000, -40, 0.15, -30]$. The stimulus protocol is a step current stimulus, where the amplitude is $\frac{0.13}{S_{soma}}$ pA/cm^2^, where we set $S_{soma}=8.32\times{10}^{-5}$ cm^2^. The duration of the step current is $1$ s and the stimulus onset is $0.4$ s. The stimulus time step is set as $0.2$ ms in the final synthetic data set. Synthetic model responses are discarded from training if their firing rates are 0, or they show spontaneous firing before stimulus onset. The discarding ratio is around 7.08.

**The stomatogastric ganglion microcircuit model of the *Cancer Borealis***

The model consists of 3 single-compartment neurons: AB/PD neuron, LP neuron, and PD neuron, where the electronically coupled AB and PD neurons are modeled as a single neuron. Each neuron has 8 kinds of currents: a Na^+^ current $I_{Na}$, a fast and a slow transient Ca^2+^ current $I_{CaT}$ and $I_{CaS}$, a transient K^+^ current $I_{A}$, a Ca^2+^-dependent K^+^ current $I_{KCa}$, a delayed rectifier K^+^ current $I_{Kd}$, a hyperpolarization-activated inward current $I_{H}$, a leak current $I_{L}$, and synaptic currents $I_{s}$. The membrane potential is modeled as Equation S9, and the synaptic current is modeled as Equation S10-S13 (7-9).

$$\begin{aligned} c_{m}\frac{dV}{dt}=-\sum_{j=1}^{N_{ion}} g_{j}\left( V-E_{j} \right)-\sum_{j=1}^{N_{synapse}} \bar{g}_{j}s_{j}\left( V-E_{j} \right)-g_{L}\left( V-E_{L} \right)+I_{noise}\#(S9) \end{aligned}$$

$$\begin{aligned} I_{s}=\bar{g}_{s}s\left( V_{post}-E_{s} \right)\#(S10) \end{aligned}$$

$$\begin{aligned} \frac{ds}{dt}=\frac{\bar{s}\left( V_{pre} \right)-s}{\tau_{s}}\#(S11) \end{aligned}$$

$$\begin{aligned} \bar{s}\left( V_{pre} \right)=\frac{1}{1+\exp\left( \frac{V_{th}-V_{pre}}{\sigma} \right)}\#(S12) \end{aligned}$$

$$\begin{aligned} \tau_{s}=\frac{1-\bar{s}(V_{pre})}{k_{-}}\#(S13) \end{aligned}$$

where $V_{post}$ and $V_{pre}$ are the membrane potential of post- and pre-synaptic neurons respectively, $V_{th}=-35$ mV is the half-activation voltage of the synapse, $\sigma=5$ mV sets the slope of the activation curve, $k_{-}$ is the rate constant for transmitter receptor dissociation rate which is $1/40$ ms for all glutamate synapses and $1/100$ ms for all cholinergic synapses, and $E_{s}=-70$ mV is the synaptic reversal potential for all glutamate synapses and $E_{s}=-80$ mV for all cholinergic synapses. At each time step, each neuron receives Gaussian noise with a mean of zero and a standard deviation of 0.001. The biophysical properties to be inferred are the maximal conductance of ion channels of each neuron: $\bar{g}_{Na}$, $\bar{g}_{CaT}$, $\bar{g}_{CaS}$, $\bar{g}_{A}$, $\bar{g}_{KCa}$, $\bar{g}_{Kd}$, $\bar{g}_{H}$ and $g_{L}$and the maximal conductances of 7 synapses: $\bar{g}_{glutamate}$and $\bar{g}_{cholinergic}$. The maximal conductance of ion channels is uniformly sampled. The lower bounds for the AB/PD neuron are: $\left[ 0, 0, 0, 0, 0, 25, 0, 0 \right]$ ms/cm^2^, and upper bounds are: $[500, 7.5, 8, 60, 15, 150, 0.2, 0.01]$ ms/cm^2^. The lower bounds for the LP neuron are: $[0, 0, 2, 10, 0, 0, 0, 0.01]$ ms/cm^2^, and upper bounds are: $[200, 2.5, 12, 60, 10, 120, 0.06, 0.04]$ ms/cm^2^. The lower bounds for the PY neuron are: $[0, 0, 0, 30, 0, 50, 0, 0]$ ms/cm^2^, and upper bounds are: $[600, 12.5, 4, 60, 5, 150, 0.06, 0.04]$ ms/cm^2^. Maximal conductances of synapses are uniformly sampled in the logarithmic domain. The lower bound and upper bound of maximal conductance of the synapse from the AB/PD neuron to the LP neuron are $0.01$ and $10000$ nS, respectively. The lower bound and upper bound of the maximal conductance of other synapses are $0.01$ and $1000$ nS, respectively. All these settings are set the same as Gonçalves et al. (9).

**Training configurations to infer biophysical properties**

The batch size is 50. The initial learning rate is 0.0001 and adapted by the Adam optimizer (10) with $\beta_{1}=0.9$ and $\beta_{2}=0.999$. The size of the similar pool is 50. The domain classifier consists of 5 fully connected layers, and the dimensions of the layers are 512, 256, 256, 128, and 128. The discrepancy loss factor is 0.0001 for the single neuron inference and 0.001 for the microcircuit inference.

We constructed a similar pool for each experimental datum, which contains the top-N most similar synthetic data to the experimental datum based on the MAE between biophysical patterns of experimental and synthetic data (**Fig. S8c & d**). Domain adversarial training (11) is used to assimilate the feature distributions of the experimental datum and its similar pool. A domain classifier $h$ is trained to classify the features as synthetic or experimental features, while the temporal feature extractor is trained to fool the domain classifier by learning domain-invariant features. The discrepancy loss is given by Equation S14-S15:

$$\begin{aligned} L_{dis}\left( f\left( D_{syn} \right),f\left( D_{exp} \right) \right)=-\frac{1}{N_{exp}} \\ \sum_{x_{exp}\in D_{exp}} \left( \mathcal{L}_{CE}\left( h\left( f\left( x_{exp} \right) \right), 1 \right)+\frac{1}{\left| SP\left( D_{syn}, x_{exp} \right) \right|}\sum_{x_{syn}\in SP\left( D_{syn}, x_{exp} \right)} \mathcal{L}_{CE}\left( h\left( f\left( x_{syn} \right) \right),0 \right) \right)\#(S14) \end{aligned}$$

$$\begin{aligned} \hat{\theta}_{h}=\arg\min_{\theta_{h}} -L_{dis}\left( f\left( D_{syn} \right),f\left( D_{exp} \right) \right)\#(S15) \end{aligned}$$

where $SP\left( D_{syn}, x_{exp} \right)$ is the similar pool of experimental data $x_{exp}$ in the synthetic dataset, and $\left| SP\left( D_{syn}, x_{exp} \right) \right|$ is the size of the similar pool, and $\theta_{h}$ is the weights of the domain classifier, and $\mathcal{L}_{CE}$ is the cross-entropy loss. At each training step, the similar pool will be updated by a new batch of synthetic data from the synthetic dataset and the synthetic data generated from the new inferences on the experimental data at the previous training step.

Synthetic data in the similar pool will be ranked based on their similarity to the corresponding experimental datum, and the synthetic label of the one ranked 1^st^ (i.e., the most similar one) will be the pseudo label of the experimental datum (**Fig. S8c**). Then self-training can be applied to the experimental data and their corresponding pseudo labels, given by Equation 4. After each training step, similar pools and pseudo labels will be updated by the new inferences of experimental data based on the similarity metric (**Fig. S8b**).

**Similarity metric to infer biophysical properties of single neurons**

We used 13 biophysical patterns in the similarity metric, which are similar to those used in Gouwens et al. (12). They are the average firing rate, action potential peak, fast trough depth, slow trough depth, time of slow trough, action potential width at half height, the latency to the first spike, the duration of the first interspike interval (ISI), the coefficient of variation of the ISIs, the average ISI, the adaptation index and the resting potential and whether there is spontaneous firing without stimulus. The similarity is calculated as the negative mean absolute error of all biophysical patterns, given by Equation S16:

$$\begin{aligned} sim\left( x_{syn}, x_{exp} \right)=-\sum_{i} \frac{\left| p_{syn}^{i}-p_{exp}^{i} \right|}{\sigma^{i}} \#(S16) \end{aligned}$$

where $x_{syn}$ and $x_{exp}$ are the synthetic data and experimental data, $p_{syn}^{i}$ and $p_{exp}^{i}$ are the $i^{\mathrm{th}}$ biophysical pattern, $\sigma^{i}$ is the smallest tolerance of the $i^{\mathrm{th}}$ pattern. The smallest tolerances of the first 12 patterns are set to be the same as in Gouwens et al. (12), and the smallest tolerance of the existence of spontaneous firing is set to be 0.001 (**Table S3**). Also, experimental data whose similarity with their corresponding synthetic samples in the similar pool is less than 6.0 will be used for domain adaptation and self-training.

**Comparison with the evolutionary search method on the inference of biophysical properties of single neurons**

The single neuronal response data of mice at a step current of amplitude 130 pA are obtained from the Allen Cell Types Database (13). Data that don’t have spikes during the stimulus window are discarded and 1052 data are used in total. For the ES method, we choose $\left( \mu+\lambda\right)$ as the evolutionary strategy (14). The probability of mating two individuals is 0.7 and the probability of mutating an individual is 0.3. The population size and offspring size are both 1000. The size of synthetic simulations of our framework is calculated as $\left( 7.08\times50+1052 \right)t$, where 7.08 is the discarding ratio when sampling the synthetic data, 50 is the batch size, and $t$ is the number of training iterations. For the comparison in **Fig. S9**, the scaling factor of discrepancy loss is 0.0001, and the self-training threshold ($S_{ST}$) and domain adaptation threshold ($S_{DA}$) are both 3. After each training iteration, inferences will be made on all experimental data and then similarity values will be calculated between each new inference and all experimental data.

**Similarity metric to infer biophysical properties of the microcircuit**

The 18 biophysical properties for the similarity calculation are adopted from Gonçalves et al. and Prinz et al. (8*,* 9): cycle period $T$, AB/PD burst duration $d_{AB}^{b}$, LP burst duration $d_{LP}^{b}$, PY burst duration $d_{PY}^{b}$, gap AB/PD end to LP start ${\Delta t}_{AB-LP}^{es}$, gap LP end to PY start ${\Delta t}_{LP-PY}^{es}$, delay AB/PD start to LP start ${\Delta t}_{AB-LP}^{ss}$, delay LP start to PY start ${\Delta t}_{LP-PY}^{ss}$, AB/PD duty cycle $d_{AB}$, LP duty cycle $d_{LP}$, PY duty cycle $d_{PY}$, phase gap AB/PD end to LP start ${\Delta\emptyset}_{AB-LP}$, phase gap LP end to PY start ${\Delta\emptyset}_{LP-PY}$, LP start phase $\emptyset_{LP}$, and PY start phase $\emptyset_{PY}$, and the maximal duration of membrane potential above $-30$ mV for each neuron ($\Delta D_{AB}$, $\Delta D_{LP}$, $\Delta D_{PY}$). If the maximal duration is below $5$ ms, it is set to $5$ ms. Some features can only be calculated if there is a rhythmic burst firing and are set to infinity if absent. The similarity metric is the same as described in Equation S16 and is calculated as the average of 10 simulations with different random seeds. The smallest tolerances of all features are summarized in **Table S4**, which are the standard deviations of features from a set of model responses generated from uniformly sampled parameters. Also, experimental data whose similarity with their corresponding synthetic samples in the similar pool is less than 0.1 will be used for domain adaptation and self-training.

**Comparison with the SNPE-C on the inference of biophysical properties of the microcircuit**

We used extracellular recordings of the microcircuit collected at $11 ^{\circ}C$, see Haddad and Marder et al. (15) for experimental details. The size of the synthetic dataset is 174000, all of which contain bursts for all neurons. 6 randomly sampled traces with a length of 12s from the extracellular recordings are used as the experimental data. We used pretrained SNPE-C from Gonçalves et al. (9), which can be found at <https://github.com/mackelab/delfi/>. The density estimator is masked autoregressive flow (16) with 5 masked autoencoders for distribution estimation (17) with [200, 400] hidden units each. There are 5 repetitions. For each repetition, we drew 100000 samples from the estimated posterior distribution and selected the one with the highest probability to calculate the MAE of biophysical patterns. The MAE reported in **Fig. S10** was the average of the 5 repetitions.

**Training software environments and hardware**

Throughout the study, we used Python 3.7 and CUDA 11. For hardware, we used NVIDIA A100 server cluster for all the training in this study.

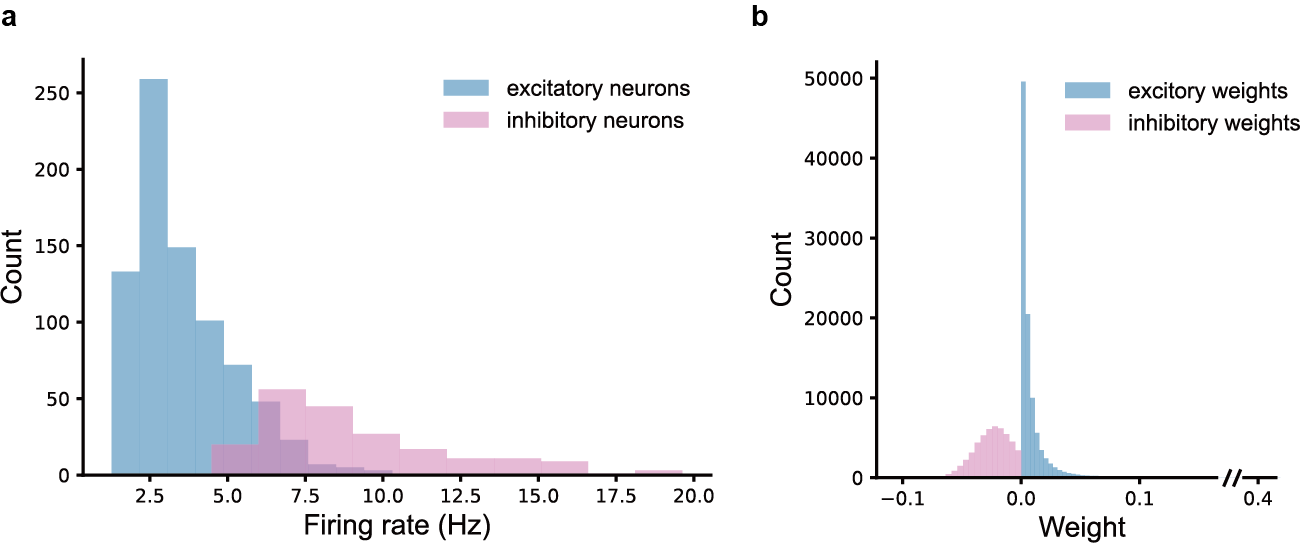

**Fig. S1. MAT network statistics. (a)** Firing rates distributions of excitatory and inhibitory neurons in the MAT network. **(b)** Synaptic weights distributions of excitatory and inhibitory synapses.

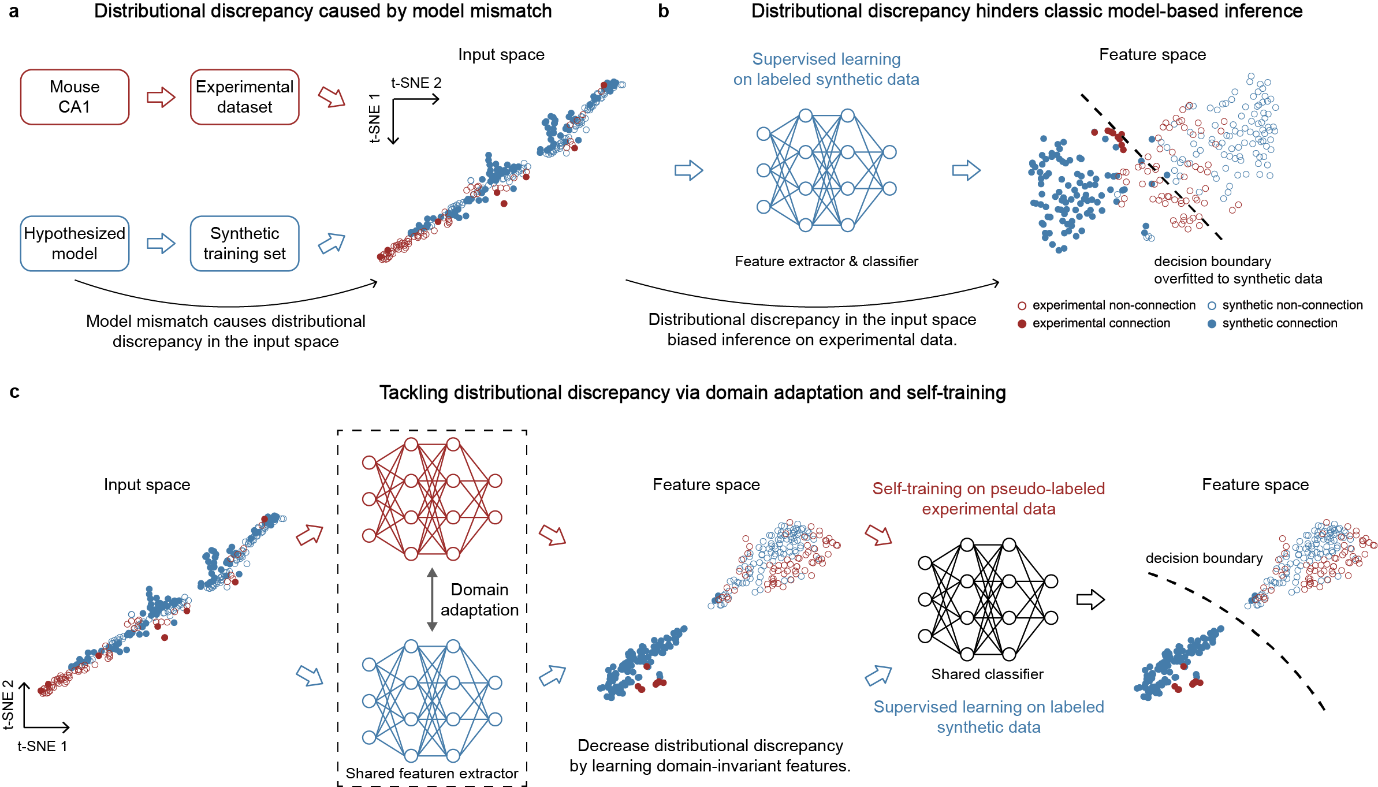

**Fig. S2. Distributional discrepancy caused by model mismatch and its solution. (a)** Projection of experimental and synthetic CCGs to the 2D space via t-SNE (18). **(b)** Traditional model-based inference framework is trained solely on labeled synthetic data, so the distributional discrepancy in the input space causes discrepancy in the feature space thus biasing the inference. **(c)** Our DeepDAM framework tackles distributional discrepancy via domain adaptivation and self-training.

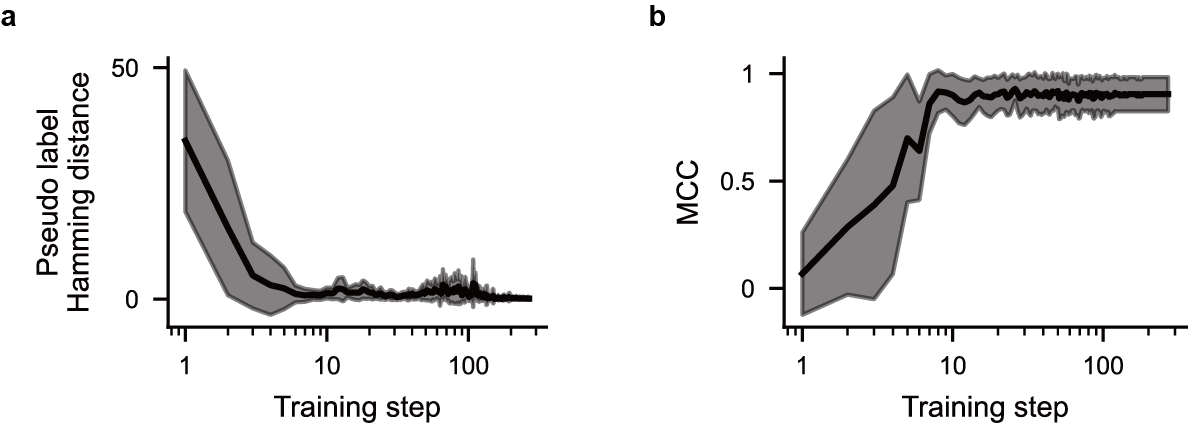

**Fig. S3. Convergence of DeepDAM. (a)** The Hamming distance between the pseudo labels at each training step and the pseudo labels at the final step. **(b)** The MCC on the experimental dataset throughout the training process.

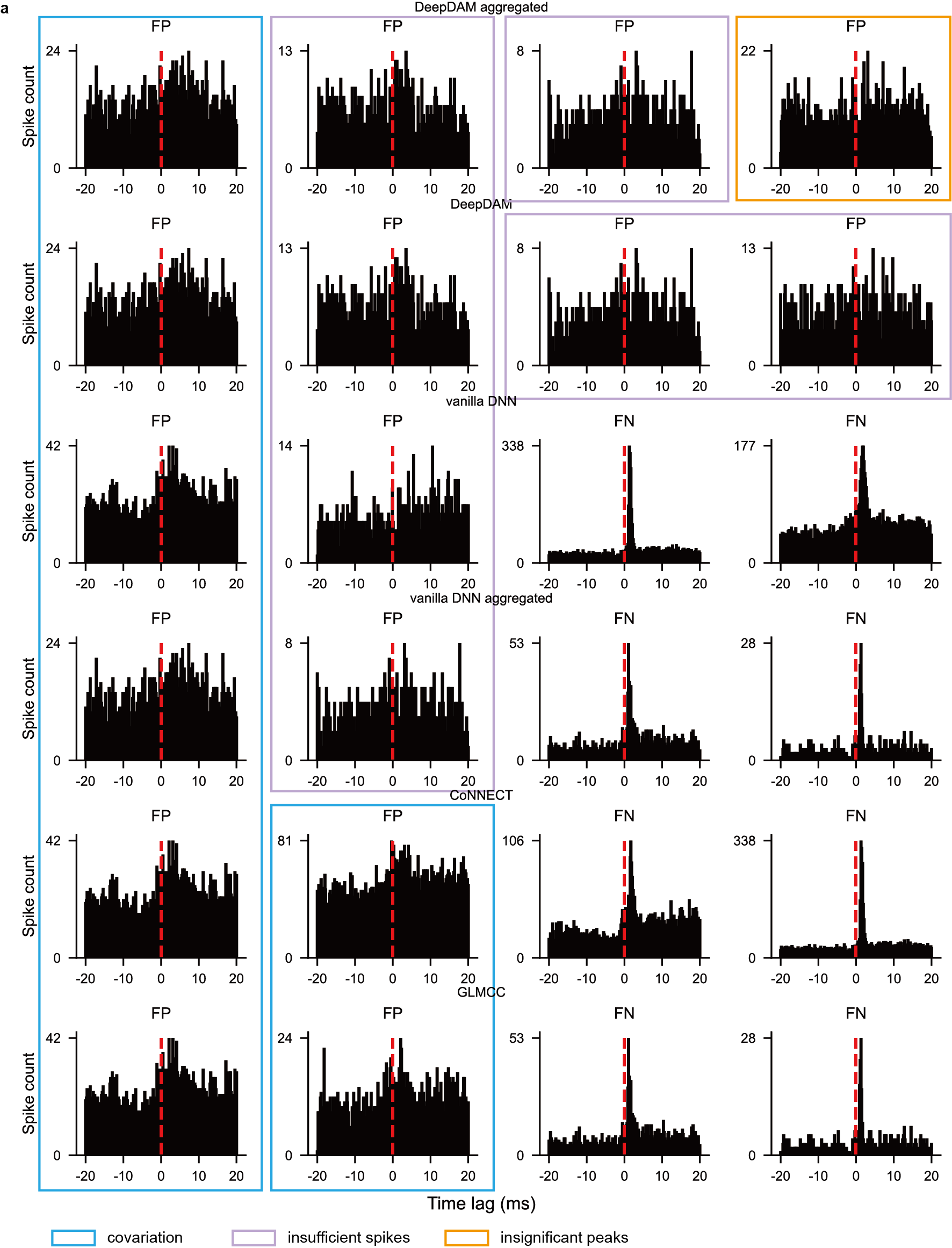

**Fig. S4. Failure cases of all methods.** (**a**) Each row is the representative failure cases of one method. FP: false positive; FN: false negative. Colored rectangular shows the reason for the false positives.

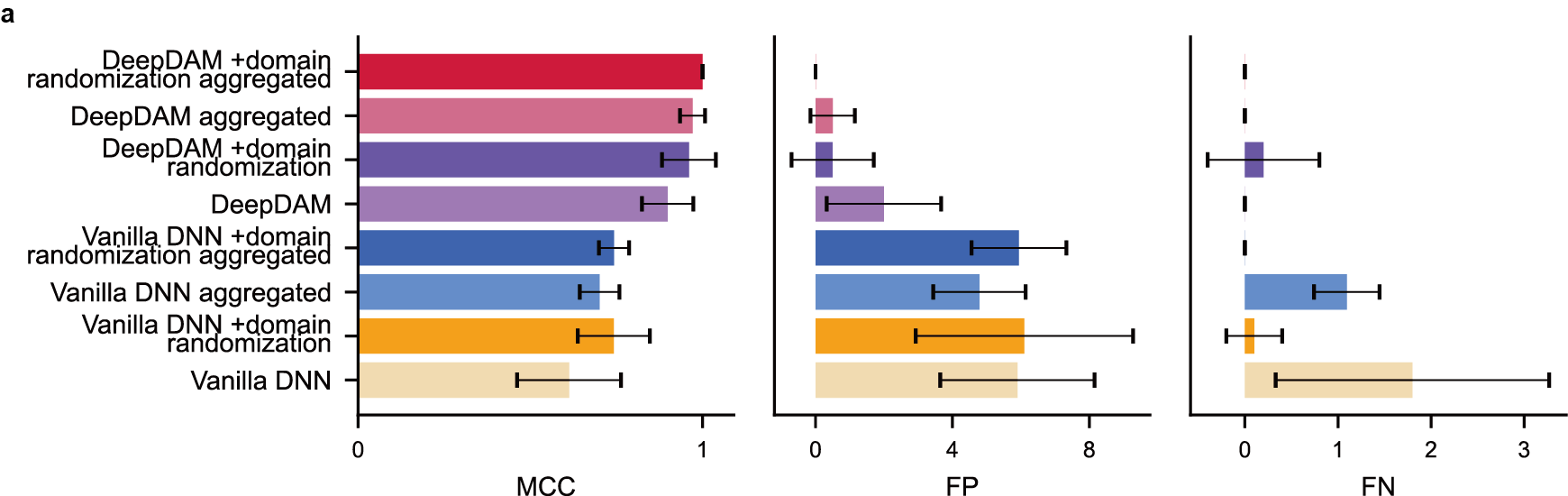

**Fig. S5. Performance comparison with domain randomization.** (**a**) Performance of DeepDAM, vanilla deep learning and their corresponding aggregated versions with and without domain randomization. MCC: Matthews correlation coefficient; FP: false positive; FN: false negative.

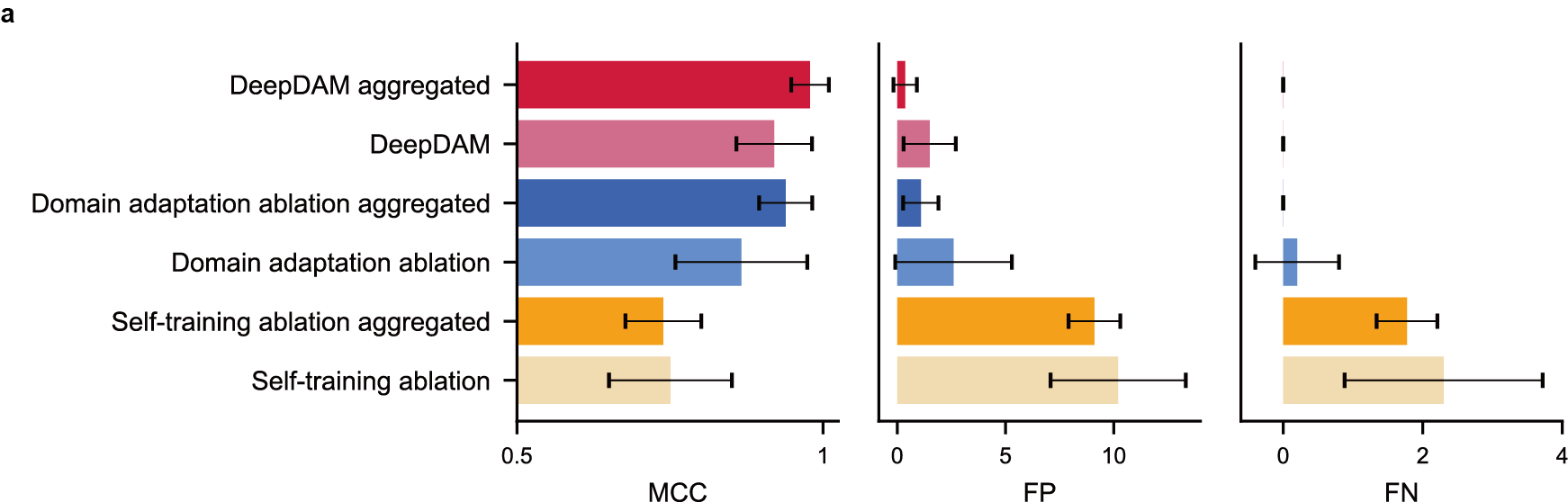

**Fig. S6. Ablation analysis of DeepDAM.** (**a**) Performance of DeepDAM, domain-adaptation ablated DeepDAM, self-training ablated DeepDAM and their corresponding aggregated versions. MCC: Matthews correlation coefficient; FP: false positive; FN: false negative.

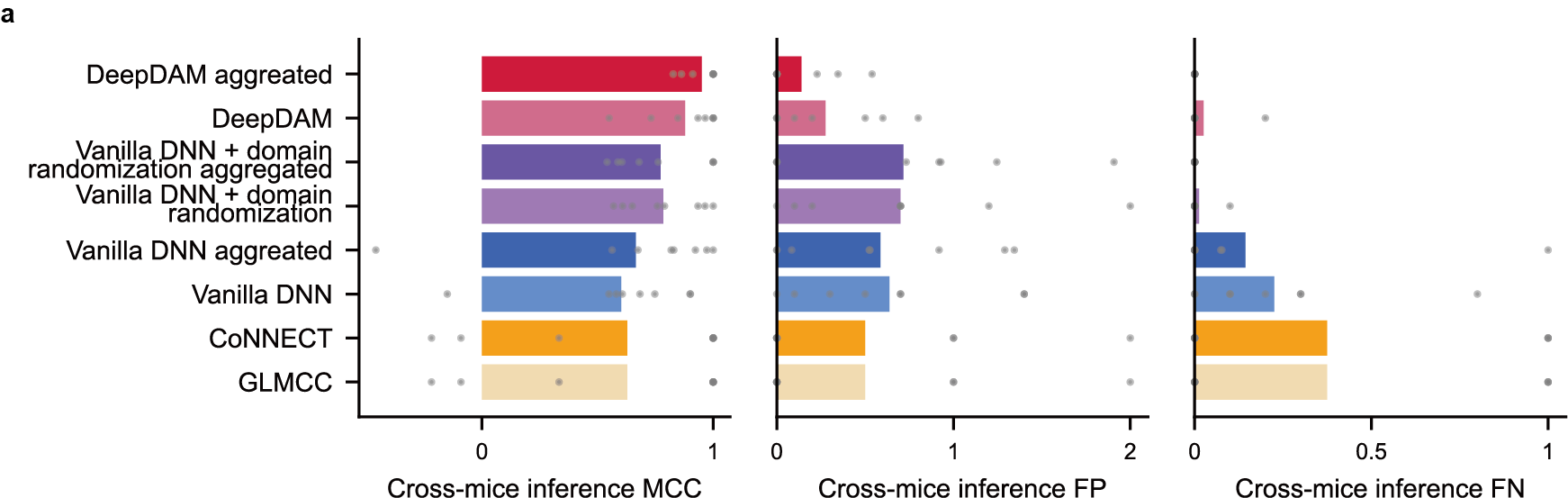

**Fig. S7. Performance of cross-mice connectivity inference.** (**a**) For DeepDAM and its aggregated version, we performed our method on N-1 (N=8) mice and evaluated the trained model on the left-out mouse. Each dot represents the performance of a mouse. We excluded two mice for this analysis as the all ground-truth labels are non-connection and the MCC cannot be computed.

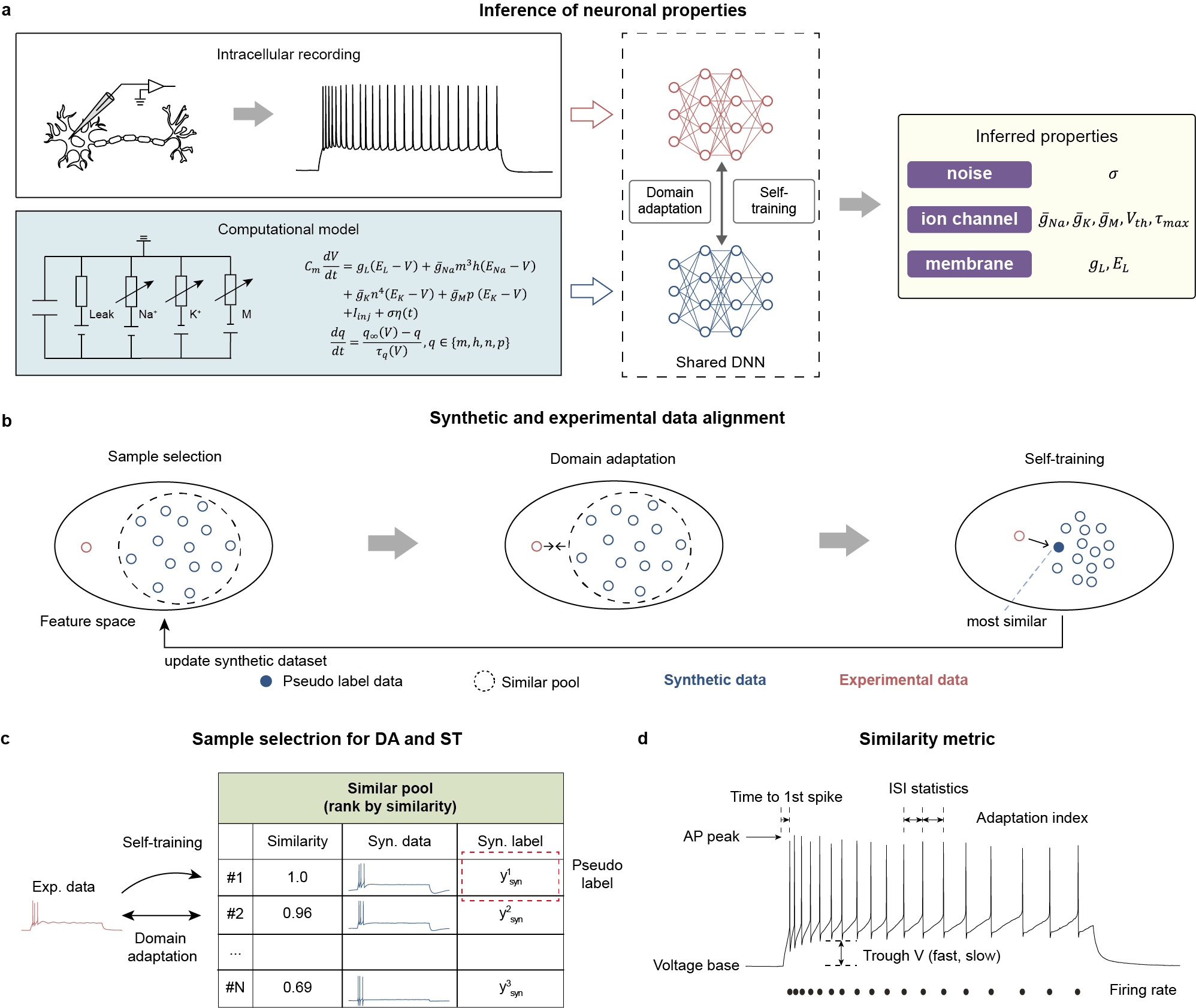

**Fig. S8. Framework instantiation to infer biophysical properties.** (**a**) The experimental data, computational model, and the biophysical properties to be inferred at the neuron scale. (**b**) Illustration of the instantiation of the DeepDAM framework to infer biophysical properties. Red symbols represent features of experimental data (exp. data) and blue ones for synthetic data (syn. data). (**c**) An illustration of the similar pool for an example experimental datum. The synthetic label of the synthetic datum that is most similar to the respective experimental datum is its pseudo label. The similarity is calculated between the synthetic model responses and their corresponding experimental data based on the similarity metric in **d**. The similarity value is normalized by the maximum similarity value. (**d**) The similarity metric for the calculation of similarities in **c** (Methods).

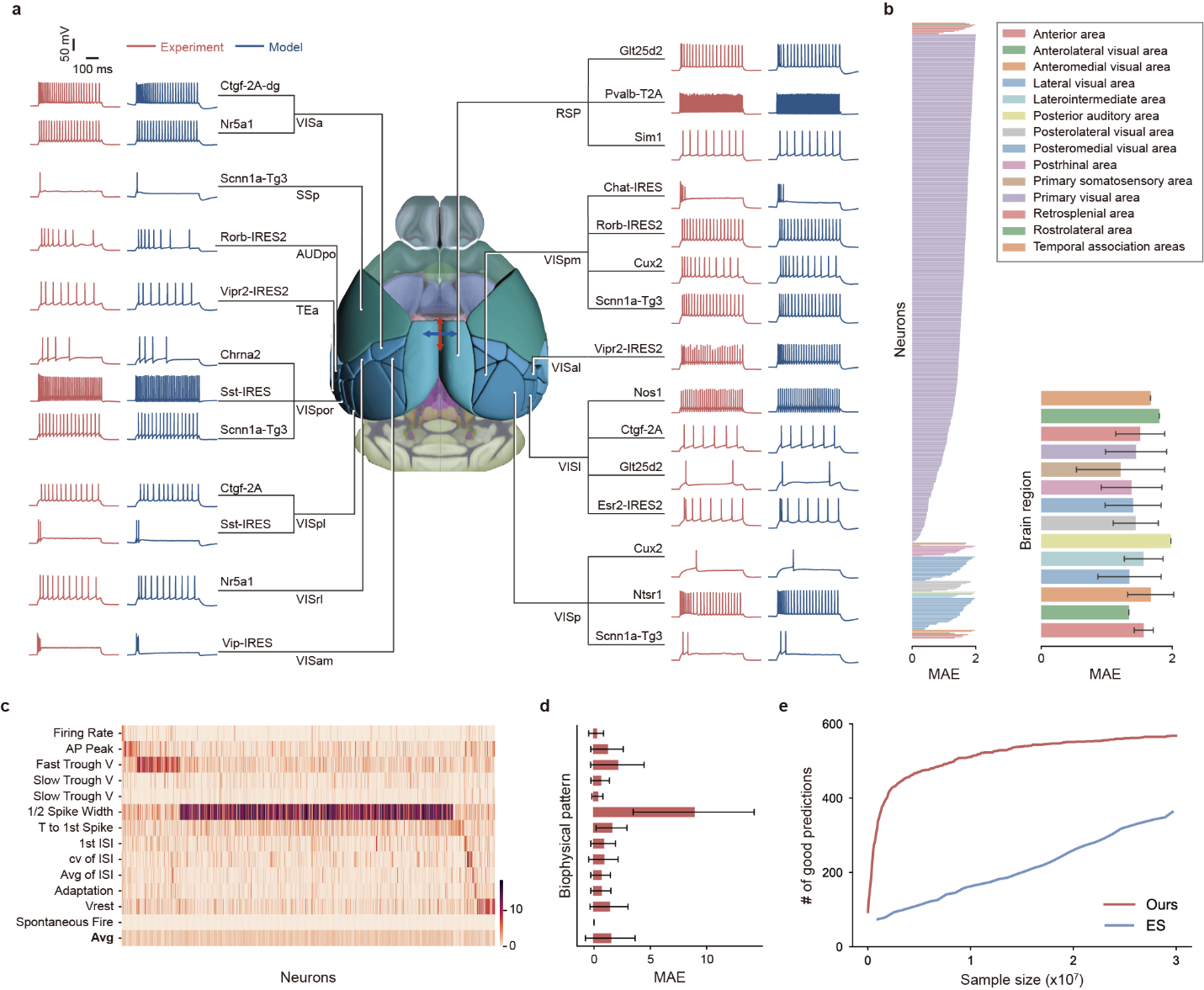

**Fig. S9. Inferring biophysical properties of neurons of mice.** (**a**) Examples of inferred results grouped by brain regions and neuron types. The brain region atlas is obtained from the Allen 3D Brain Explorer (19). Blue curves: the model responses are generated by applying the same stimulus protocol with the inferred properties. Red curves: experimental data taken from Allen Cell Types Database (13). (**b**) Mean absolute error (MAE) of biophysical patterns between the model responses and the experimental data of 574 neurons sorted by brain regions. Insect: the average MAE of all neurons in the brain region. Error bar: standard deviation. (**c**) MAE of each biophysical pattern of 574 neuron models, sorted by the biophysical patterns with the maximum error. (**d**) MAE of each feature across all neuron models in **c**. The y-axis has the same order of labels as in c. Error bar: standard deviation across 574 models. (**e**) Sample efficiency compared to the evolutionary search (ES) method (9). The y-axis is the number of good predictions from the two methods, where good is defined as predictions whose MAE of biophysical patterns is smaller than 2.0.

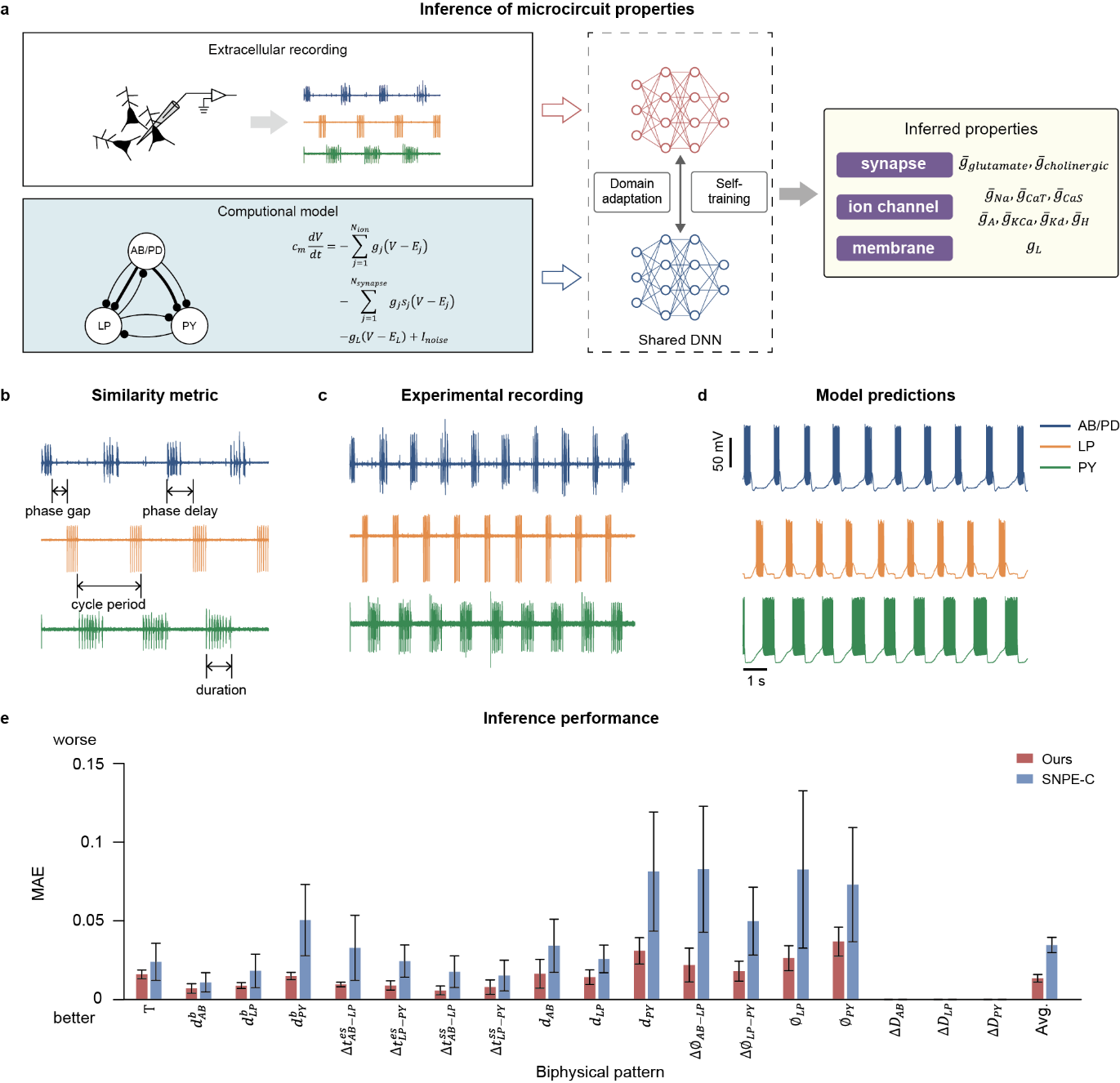

**Fig. S10. Inferring biophysical properties of a microcircuit of the *Cancer Borealis*.** (**a**) The experimental data, computational model, and the neural properties to be inferred at the microcircuit scale. In the illustration of the computational model panel, the thin connections are fast glutamatergic synapses, and the thick connections are slow cholinergic synapses. (**b**) The similarity metric for this task. Different pyloric statistics are compared for the calculation of similarity. (**c**) The experimental extracellular recordings of the microcircuit. (**d**) The predicted model responses by applying a similar stimulus protocol as the experiment based on a set of inferred properties of the model. (**e**) Performance comparison with SNPE-C (9) of MAE of biophysical patterns. The performance of SNPE-C was given by the pretrained model from Gonçalves et al (9) (Methods). Error bar: standard deviation across 5 random repetitions.

**Table S1. Sensitivity analysis of hyperparameters of DeepDAM.**

| $\boldsymbol{\lambda}_{\boldsymbol{dis}}$ | $\boldsymbol{G}_{\boldsymbol{DA}}$ | $\boldsymbol{G}_{\boldsymbol{ST}}$ | Aggregated MCC mean±std | Original MCC mean±std | Aggregated FP mean±std | Original FP mean±std | Original FN mean±std | Original FN mean±std |
| --- | --- | --- | --- | --- | --- | --- | --- | --- |
| 0.01 | 0.05 | 0.05 | 0.91±0.06 | 0.85±0.12 | 1.80±1.21 | 3.5±3.04 | 0.0±0.0 | 0.0±0.0 |
| 0.01 | 0.05 | 0.01 | 0.91±0.06 | 0.85±0.12 | 1.80±1.21 | 3.5±3.04 | 0.0±0.0 | 0.0±0.0 |
| 0.01 | 0.05 | 0.001 | 0.91±0.06 | 0.85±0.12 | 1.80±1.21 | 3.5±3.04 | 0.0±0.0 | 0.0±0.0 |
| 0.01 | 0.01 | 0.01 | 0.97±0.04 | 0.91±0.06 | 0.59±0.64 | 1.6±1.11 | 0.0±0.0 | 0.0±0.0 |
| 0.01 | 0.01 | 0.001 | 0.97±0.04 | 0.92±0.06 | 0.56±0.65 | 1.6±1.2 | 0.0±0.0 | 0.0±0.0 |
| 0.01 | 0.001 | 0.001 | 0.68±0.06 | 0.63±0.10 | 5.69±1.29 | 6.6±1.69 | 0.93±0.60 | 1.3±1.27 |
| 0.001 | 0.05 | 0.05 | 0.91±0.05 | 0.86±0.11 | 1.73±1.05 | 3.0±2.79 | 0.0±0.0 | 0.1±0.3 |
| 0.001 | 0.05 | 0.01 | 0.91±0.05 | 0.85±0.11 | 1.73±1.05 | 3.1±2.74 | 0.0±0.0 | 0.1±0.3 |
| 0.001 | 0.05 | 0.001 | 0.91±0.05 | 0.85±0.11 | 1.73±1.05 | 3.1±2.74 | 0.0±0.0 | 0.1±0.3 |
| 0.001 | 0.01 | 0.01 | 0.97±0.04 | 0.91±0.09 | 0.49±0.65 | 1.8±2.04 | 0.0±0.0 | 0.0±0.0 |
| 0.001 | 0.01 | 0.001 | 0.97±0.04 | 0.91±0.09 | 0.56±0.69 | 1.9±2.02 | 0.0±0.0 | 0.0±0.0 |
| 0.001 | 0.001 | 0.001 | 0.69±0.06 | 0.63±0.12 | 5.65±1.28 | 6.5±1.86 | 0.92±0.60 | 1.3±1.35 |
| 0.0001 | 0.05 | 0.05 | 0.92±0.04 | 0.86±0.11 | 1.39±0.86 | 2.8±2.79 | 0.0±0.0 | 0.2±0.6 |
| 0.0001 | 0.05 | 0.01 | 0.92±0.04 | 0.86±0.11 | 1.39±0.86 | 2.8±2.79 | 0.0±0.0 | 0.2±0.6 |
| 0.0001 | 0.05 | 0.001 | 0.94±0.04 | 0.87±0.11 | 1.09±0.82 | 2.6±2.69 | 0.0±0.0 | 0.2±0.6 |
| 0.0001 | 0.01 | 0.01 | 0.97±0.04 | 0.91±0.08 | 0.53±0.74 | 1.9±1.81 | 0.0±0.0 | 0.0±0.0 |
| 0.0001 | 0.01 | 0.001 | 0.98±0.03 | 0.91±0.08 | 0.34±0.58 | 1.8±1.83 | 0.0±0.0 | 0.0±0.0 |
| 0.0001 | 0.001 | 0.001 | 0.68±0.06 | 0.63±0.10 | 5.71±1.25 | 6.6±1.86 | 0.93±0.60 | 1.3±1.27 |
| Vanilla deep learning | | | 0.70±0.06 | 0.61±0.15 | 4.79±1.35 | 5.9±2.26 | 1.1±0.35 | 1.8±1.47 |

**Table S2. Performance with domain randomization.**

| $\boldsymbol{\lambda}_{\boldsymbol{dis}}$ | $\boldsymbol{G}_{\boldsymbol{DA}}$ | $\boldsymbol{G}_{\boldsymbol{ST}}$ | Aggregated MCC mean±std | Original MCC mean±std | Aggregated FP mean±std | Original FP mean±std | Original FN mean±std | Original FN mean±std |
| --- | --- | --- | --- | --- | --- | --- | --- | --- |
| 0.01 | 0.05 | 0.05 | 1.0±0.0 | 0.96±0.08 | 0.0±0.0 | 0.5±1.20 | 0.0±0.0 | 0.2±0.6 |
| 0.01 | 0.05 | 0.01 | 1.0±0.0 | 0.96±0.08 | 0.0±0.0 | 0.5±1.20 | 0.0±0.0 | 0.2±0.6 |
| 0.01 | 0.05 | 0.001 | 1.0±0.0 | 0.96±0.08 | 0.0±0.0 | 0.5±1.20 | 0.0±0.0 | 0.2±0.6 |
| 0.01 | 0.01 | 0.01 | 0.97±0.03 | 0.91±0.06 | 0.48±0.58 | 1.7±1.19 | 0.0±0.0 | 0.0±0.0 |
| 0.01 | 0.01 | 0.001 | 0.97±0.04 | 0.91±0.06 | 0.51±0.63 | 1.7±1.19 | 0.0±0.0 | 0.0±0.0 |
| 0.01 | 0.001 | 0.001 | 0.74±0.04 | 0.72±0.09 | 6.18±1.32 | 6.6±2.97 | 0.0±0.0 | 0.1±0.3 |
| 0.001 | 0.05 | 0.05 | 1.0±0.0 | 0.96±0.08 | 0.0±0.0 | 0.5±1.20 | 0.0±0.0 | 0.2±0.6 |
| 0.001 | 0.05 | 0.01 | 1.0±0.0 | 0.96±0.08 | 0.0±0.0 | 0.5±1.20 | 0.0±0.0 | 0.2±0.6 |
| 0.001 | 0.05 | 0.001 | 1.0±0.0 | 0.96±0.08 | 0.0±0.0 | 0.5±1.20 | 0.0±0.0 | 0.2±0.6 |
| 0.001 | 0.01 | 0.01 | 0.98±0.03 | 0.93±0.06 | 0.37±0.54 | 1.3±1.19 | 0.0±0.0 | 0.0±0.0 |
| 0.001 | 0.01 | 0.001 | 0.97±0.04 | 0.91±0.06 | 0.54±0.65 | 1.7±1.19 | 0.0±0.0 | 0.0±0.0 |
| 0.001 | 0.001 | 0.001 | 0.73±0.04 | 0.72±0.09 | 6.29±1.28 | 6.6±2.97 | 0.0±0.0 | 0.1±0.3 |
| 0.0001 | 0.05 | 0.05 | 1.0±0.0 | 0.96±0.08 | 0.0±0.0 | 0.5±1.20 | 0.0±0.0 | 0.2±0.6 |
| 0.0001 | 0.05 | 0.01 | 1.0±0.0 | 0.96±0.08 | 0.0±0.0 | 0.5±1.20 | 0.0±0.0 | 0.2±0.6 |
| 0.0001 | 0.05 | 0.001 | 1.0±0.0 | 0.96±0.08 | 0.0±0.0 | 0.5±1.20 | 0.0±0.0 | 0.2±0.6 |
| 0.0001 | 0.01 | 0.01 | 0.98±0.03 | 0.92±0.07 | 0.35±0.52 | 1.6±1.56 | 0.0±0.0 | 0.0±0.0 |
| 0.0001 | 0.01 | 0.001 | 0.96±0.04 | 0.90±0.07 | 0.66±0.68 | 2.0±1.48 | 0.0±0.0 | 0.0±0.0 |
| 0.0001 | 0.001 | 0.001 | 0.74±0.04 | 0.74±0.10 | 6.18±1.31 | 6.2±3.22 | 0.0±0.0 | 0.1±0.3 |
| Vanilla deep learning | | | 0.74±0.04 | 0.74±0.11 | 5.94±1.39 | 6.1±3.18 | 0.0±0.0 | 0.1±0.3 |

**Table S3.** **The smallest tolerances used in the similarity calculation of biophysical patterns of single neurons (adapted from** **Gouwens et al. (**12**)).**

| **Biophysical pattern** | **Smallest tolerance (**$\boldsymbol{\sigma}$**)** |
| --- | --- |
| Average firing frequency | 0.5 Hz |
| Latency to the first spike | 5 ms |
| Average interspike interval (ISI) | 0.5 ms |
| Duration of first ISI | 1 ms |
| Coefficient of variation of ISIs | 0.1 |
| Adaptation index | 0.001 |
| Action potential peak | 2 mV |
| Fast trough depth | 2 mV |
| Slow trough depth | 2 mV |
| Time of slow trough (as a fraction of ISI) | 0.05 |
| Action potential width at half-height | 0.1 ms |
| Resting potential | 2 mV |
| Average firing frequency | 0.5 Hz |
| Existence of spontaneous firing | 0.001 |

### **Table S4. The smallest tolerances used in the similarity calculation of biophysical patterns of the microcircuit.**

| **Biophysical pattern** | **Smallest tolerance (**$\boldsymbol{\sigma}$**)** |
| --- | --- |
| $T$ | 1063.169 ms |
| $d_{AB}^{b}$ | 733.209 ms |
| $d_{LP}^{b}$ | 665.553 ms |
| $d_{PY}^{b}$ | 459.875 ms |
| ${\Delta t}_{AB-LP}^{es}$ | 433.098 ms |
| ${\Delta t}_{LP-PY}^{es}$ | 705.638 ms |
| ${\Delta t}_{AB-LP}^{ss}$ | 859.753 ms |
| ${\Delta t}_{LP-PY}^{ss}$ | 909.732 ms |
| $d_{AB}$ | 0.240 ms |
| $d_{LP}$ | 0.337 ms |
| $d_{PY}$ | 0.196 ms |
| ${\Delta\emptyset}_{AB-LP}$ | 0.164 s |
| ${\Delta\emptyset}_{LP-PY}$ | 0.296 s |
| $\emptyset_{LP}$ | 0.191 s |
| $\emptyset_{PY}$ | 0.205 s |
| $\Delta D_{AB}$ | 29.397 ms |
| $\Delta D_{LP}$ | 58.110 ms |
| $\Delta D_{PY}$ | 35.565 ms |
